# PerTurboAgent: A Self-Planning Agent for Boosting Sequential Perturb-seq Experiments

**DOI:** 10.1101/2025.05.25.656020

**Authors:** Minsheng Hao, Yongju Lee, Hanchen Wang, Gabriele Scalia, Aviv Regev

**Author notes:** Work done during an internship at Genentech (Minsheng Hao).

## Abstract

Understanding how genetic interventions affect a cell’s phenotype is key to revealing causal gene regulatory mechanisms and finding novel drug targets. Pooled, high-content perturbation screening methods like Perturb-seq allow us to assess the impact of each of a large number of genetic interventions on a rich cellular profile of RNA or other features, facilitating such discoveries. However, the overall scale of perturbations, especially when considering combinations of genes for perturbation, cannot be tackled exhaustively in the lab. An alternative is to use an iterative design: By leveraging the modularity and sparsity of gene circuits along with prior biological knowledge, we can predict the impact of unseen genetic perturbations on these profiles and group genes with similar effects into co-functional modules, followed by a new round of Perturb-Seq to test these predictions and improve the overall performance of the model. These iterative cycles of experiment and prediction allow prioritizing genes for testing, maximizing the knowledge gleaned from fixed experimental resources, and opening the way to learn general predictive models. Designing these experiments requires a system that can analyze a cellular system, incorporate new and existing knowledge, use statistical tools, predict the effects of unseen perturbations, and prioritize the set of perturbations for the next iteration. These can be time-consuming tasks for scientists and require multiple different skills. Here, we developed PerTurboAgent, an LLM-based agent that excels in predicting candidate gene panels for iterative Perturb-Seq experiments through self-directed data analysis and knowledge retrieval. We evaluated PerTurboAgent based on its ability to identify genes with a phenotypic impact on gene expression upon perturbation in genome-scale perturbation data. PerTurboAgent outperforms existing agent-based and active learning strategies, offering an efficient and understandable approach to designing sequential perturbation experiments.

## 1 Introduction

Uncovering the causal genetic mechanisms underlying a cell’s phenotype is a fundamental problem in cell biology and has multiple practical implications, from the discovery of drug targets to the engineering of efficacious cell therapies with desired states. A major experimental strategy to recover causal mechanisms relies on intervention by genetic perturbation followed by measurement of the impact on the cell’s phenotype (Xia et al., 2021). In recent years, Perturb-seq, a pooled CRISPR-based genetic screen with single cell profiling readout, has emerged as a highly scaled and impactful approach, because it combines a large number of perturbations assays simultaneously with the impact of each perturbation on a complex, high-content readout, such as a single cell RNA-seq (scRNA-Seq) profile. Perturb-Seq has been applied at a genome scale to assess the impact of perturbing each of thousands of individual genes or subsets of pair-wise combinations (Dixit et al., 2016; Norman et al., 2019; Nadig et al., 2024; Replogle et al., 2022). The resulting measurements have been used both to directly assess perturbation effects, reconstruct modular models of gene regulation, and train models that attempt to predict the impact of unobserved perturbations (with partial success) (Adamson et al., 2016; Geiger-Schuller et al., 2023; Bunne et al., 2023; Miladinovic et al., 2025; Roohani et al., 2024a).

Despite these advances, the breadth of biological conditions (across cell types, states, and environments) and the number of possible perturbations (across individual genes and their multi-way combinations) make exhaustive experiments impossible. Fortunately, because gene circuits are often sparse and modular, it should, in principle, be possible to use the information drawn from perturbation experiments to train models that generalize to the full perturbation space, without exhaustively testing all perturbations in the lab (Yeung & Ruzzo, 2001; Brunet et al., 2004; Yao et al., 2024). Indeed, knowledge about correlated gene expression patterns and shared gene functions has been leveraged for machine learning models, including generative and foundation models which aim to predict the impact of unseen perturbations (Roohani et al., 2024a; Bunne et al., 2023; Cui et al., 2024; Lotfollahi et al., 2019; 2023; Hao et al., 2024). However, such models are currently only partly successful at best and ideally require iterative validation experiments to test their predictions and improve the models’ performance (Rood et al., 2024).

In this context, an intriguing challenge is to design cost-effective experiments, leveraging all recent knowledge and advanced tools. Such sequential (or iterative) experiments can strategically prioritize genes for perturbation based on their potential to either enhance the overall predictive performance of the model or how informative perturbations should be towards discovering the gene circuits underlying a specific target phenotype. In this paradigm, experimental plans are iteratively updated by the model based on newly acquired data, thereby enhancing the information gained for a certain cost compared to acquiring all data at once (Greenhill et al., 2020). Active learning and Bayesian optimization have been instrumental in developing methods for sequential designs for perturbation experiments. For instance, *GeneDisco* and *DiscoBAX* refined perturbation selection purely based on experimental outcomes, without leveraging biological knowledge (Mehrjou et al., 2021; Lyle et al., 2023). *Iterpert* advanced these by integrating prior knowledge into a kernel-based active learning framework, though its optimization still centers on enhancing prediction model performance rather than pinpointing perturbations most relevant to a specific target phenotype (Huang et al., 2024b).

More recently, large language models (LLMs) have shown promise in biomedical applications, such as interpreting literature, creating gene embeddings, and tackling tasks in Scientific Q and A and RNA design, including incorporation in an agent that can plan, execute, and revise plans to solve a given task based on the results of their previous actions (Lu et al., 2022; Huang et al., 2024a; Gao et al., 2024; Swanson et al., 2024; Lu et al., 2024). The sequential Perturb-seq design problem is well-suited for LLM agents, because of the need to integrate previous knowledge, analyze experimental results, and incorporate machine learning predictions to select candidate genes effectively. LLM agents can orchestrate such different tools as action items, reflect on the results of each action reasonably, and plan the next experiment Lee et al. (2024). Such agents can use multiple information sources, mimic expert workflows, integrate analytical actions, from gene set enrichment analysis to predictive modeling of perturbation outcomes, and dynamically adjust plans as new data emerge. A recent *BioDiscoveryAgent* employs LLMs with tools for information processing and gene selection, but relies on fixed plans and doesn’t fully leverage gene expression data Roohani et al. (2024b).

Here, we propose *PerTurboAgent*, a self-planning LLM-based agent designed to enhance sequential Perturb-seq experiments. We develop diverse actions across three categories to enable full utilization of experimental data, prior knowledge, and LLM capabilities: agent-based (prediction, reflection, refinement), data-driven (e.g., gene set enrichment tests), and prediction model-driven actions. Furthermore, we introduce an action memory to support multi-step reasoning and execution during perturbation selection. The agent’s chosen actions and results are recorded, allowing subsequent steps to adaptively evolve the plan. Experiments across eleven phenotypes demonstrate that PerTurboAgent surpasses previous methods. By analyzing action frequency and internal memory, PerTurboAgent provides interpretable insights and transparent reasoning. We show that PerTurboAgent is compatible with both closed-source and open-source models, while also benefiting from the more advanced models. Overall, PerTurboAgent transcends fixed strategies to offer a flexible, context-aware solution for identifying gene perturbations most strongly associated with target phenotypes.

## 2 Problem Setting

Given a cellular system described by a set of genes *G* = {*p*_1_, *p*_2_, …, *p*_*n*_}, we define a specific cellular function or phenotype by a subset of *k* genes, *P* = {*p*_1_, *p*_2_, …, *p*_*k*_}, termed *associated descriptive genes* (ADGs). The cellular system can be regulated through a set of genetic interventions *I* = {*i*_1_, *i*_2_, …, *i*_*h*_}. We define the *hit genes I*_*truth*_ ⊂ *I* as the set of genetic interventions that induce a stronger expression change of *P* compared to control (unperturbed) cells than the other genetic interventions, where the expression change is set based on a z-score threshold as previously defined (Replogle et al., 2022).

The goal is to identify *I*_*truth*_ through *N* sequential rounds of experiments. In each round *R*, the agent selects a set of *m* genes 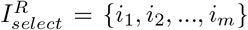 from *I* and obtains the associated experimental results. The agent can select each gene from *I* only once. We refer to all genes selected until round *R* as *I*_*tested*_, and unselected genes as *I*_*untested*_. At the initial round 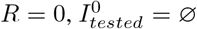 and the untested set 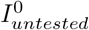 is equal to *I*. The agent can choose multiple actions among the given action pool at each round, before making the selection 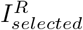. However, the round ends after the agent obtains the experimental results. We summarize the selection process as follows:

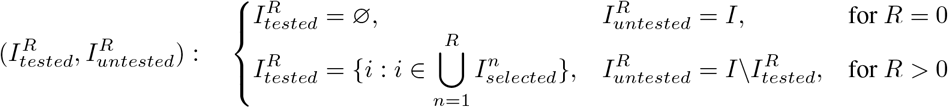

Unlike previous studies (Roohani et al., 2024b), we enable the LLM agent to understand both control and perturbed cells by providing expression profiles of both unperturbed and perturbed cells. At the initial round, the LLM agent receives the control cells’ gene expression *G*_*c*_. In each subsequent round, it obtains the gene expression profiles from perturbed cells *G*_*p*_ for the *m* selected genetic perturbation in 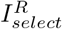. The agent directly accesses the raw gene expression data and analyzes it according to its plan.

Additionally, we introduce a *phenotype score, S*_*P*_, defined as the expression change of genes in *P* after perturbation, compared with the control sample:

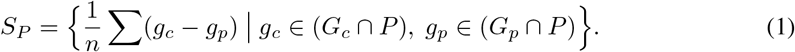

This allows the agent to assess the strength of the selected perturbations’ impact on the given phenotype *P*. These data are stored to facilitate subsequent rounds of gene selection.

## 3 PerTurboAgent

PerTurboAgent is a self-planning agent for sequential experiment workflows. At each round, the agent tackles gene selection as a multi-step decision-making process (Figure 1). At each step *K*, the agent selects an action based on the context of the target phenotype, along with all prior actions and outcomes within the current round. To achieve this, the agent is equipped with an *action memory*. After executing an action, the action-result pair is stored for reference in subsequent steps. A detailed multi-step decision-making log of PerturbAgent in a single round is provided in Section A.4.

**Figure 1:**
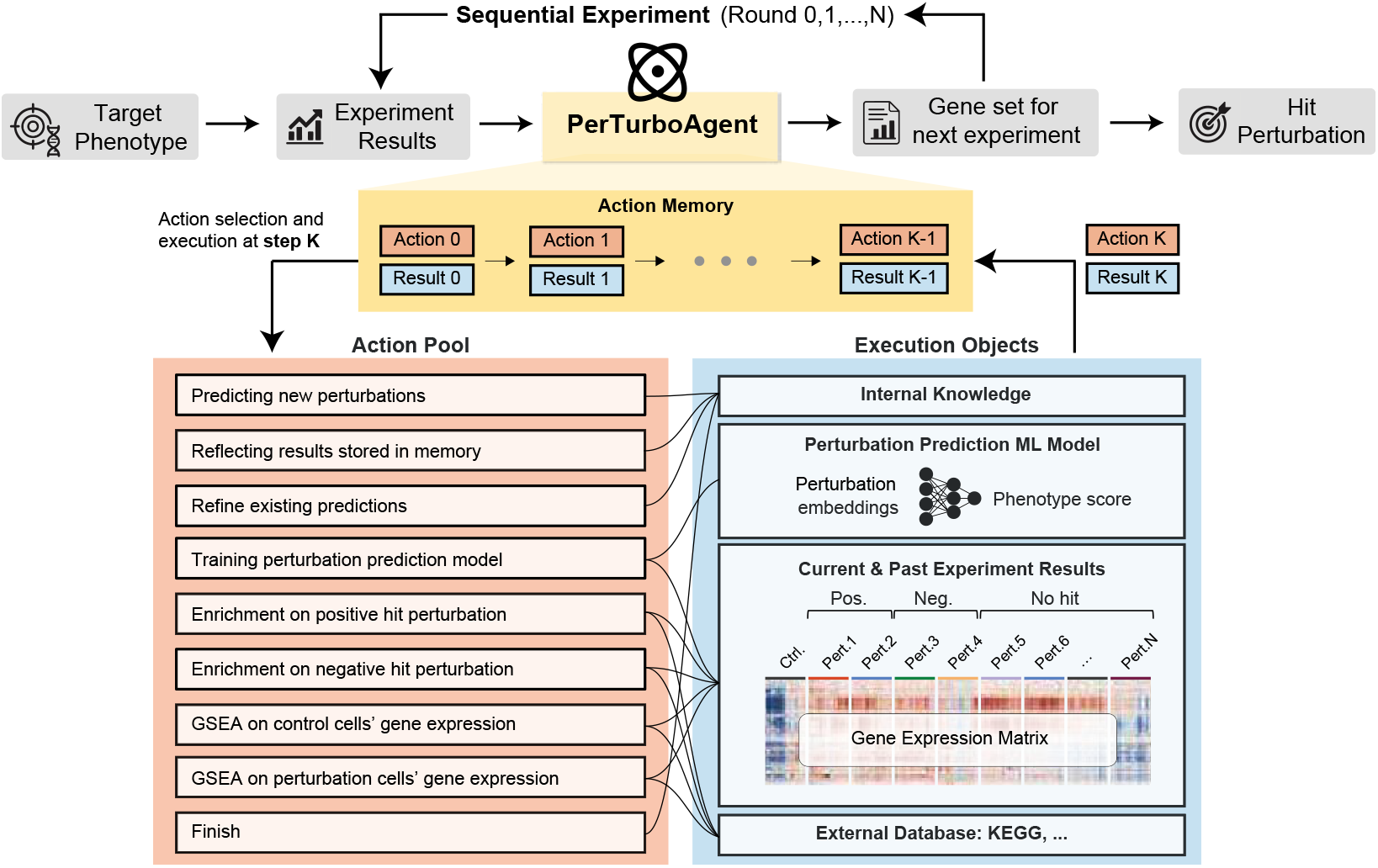
PerTurboAgent overview. PerTurboAgent sequentially identifies gene sets with key regulatory impact (Hit Perturbations) on a target phenotype. In each experimental round, PerTurboAgent selects a limited number of perturbation experiments to maximize the number of identified Hit Perturbations by the final round. At each round, PerTurboAgent takes actions from its Action Pool, iteratively choosing multiple actions. The outcomes of these actions are stored in Action Memory, a memory bank for each round, enabling PerTurboAgent to make informed decisions based on past actions and results.

### 3.1 Task Initialization

At the beginning of each round, the agent is initialized with a task definition prompt and an action-loading prompt. The task definition prompt specifies the target phenotype name and the functional gene set related to that phenotype (associated descriptive genes (ADGs)), as defined by the user as part of the phenotype *P* (*e*.*g*., translation, glycolysis, growth signaling, etc). A user can define ADGs based on prior knowledge and curation, or by using the results of a Perturb-Seq experiment in the same or another system to choose a phenotype of interest (e.g., *P* : Translation, *ADGs*: ‘ACTG1’, ‘CCNG1’, ‘COX7C’, ‘EEF2’, ‘EIF2S3’, ‘EIF3F’). The action-loading prompt provides the names and descriptions of all available actions. The agent selects an action based on its assigned code number and executes it to obtain the final results (Section A.1).

### 3.2 Action Pool

All actions in the pool are categorized into one of three groups based on their execution objectives: *Reasoning, ML Inference*, or *Analysis*.

*Reasoning* actions leverage the agent’s own capabilities to predict new perturbations, reflect on results, and refine predictions (Section A.2.1):

- *Selecting new perturbations*. The agent predicts the gene symbols to populate the perturbation sets 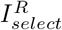 in round *R*. The Agent responses are structured into three parts: *Reflection, Research Plan*, and *Solution* to enhance interpretability. Because LLM hallucinations can lead the Agent to provide non-existent gene names, we require the agent to match the predicted genes in the *Solution* against a gene name database, thus distinguishing a valid and invalid set in the results.
- *Reflecting on stored results*. The agent reviews prior results stored in its memory and analyzes previous steps.
- *Refining existing predictions*. The agent critiques and updates its predictions, removing unreasonable perturbations and adding new ones. Responses are structured into three parts: *Critique, SolutionRemoval*, and *SolutionAddition*.
- *Finish*. This action allows the agent to terminate the loop and return its final set of predictions.

*ML Inference* actions focus on training models to prioritize perturbations based on their predicted phenotype scores (Section A.2.2):

- *Training ML model for perturbation prediction*. At round T, the agent invokes functions to train a LightGBM regressor Ke et al. (2017), which is a light ML model that is easily trainable between each round, using the GenePT embeddings Chen & Zou (2024) of all previously tested perturbations 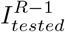. The model is trained to predict phenotype scores *S*_*P*_ based on experimental results observed until previous rounds. After training, the model is used to estimate *S*_*P*_ for untested perturbations 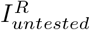, and selects a predefined number of top-scoring perturbations 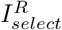.
- *Delayed activation of ML model*. In early rounds, when there is very little training data, the results from the regressor would mislead the agent. Thus, we allow the agent to choose the ML training action after seeing the increment of hits from two consecutive rounds. We ablated the effect of this heuristic design in Section 5.3.

*Analysis* actions analyze accumulated gene expression data to extract insights (Section A.2.3):

- *Enrichment of positive/negative hit perturbations*. Given a gene set containing all positive/negative hits, the agent performs enrichment analysis to test whether these hits are significantly enriched for genes from predefined pathways or cellular processes. The agent selects the enrichment reference database by itself and identifies significant pathways.
- *GSEA on gene expression from control or perturbed cells*. Using Gene Set Enrichment Analysis (GSEA), the agent identifies significantly enriched pathways based on gene expression profiles from control or newly perturbed cells. Unlike the previous enrichment analysis, which finds associations at the gene set level, GSEA directly analyzes gene expression data to uncover enriched pathways for each perturbation.

We set a maximum of 20 action steps per round, and the agent is aware of the current step index. At each step, the agent selects a single action. The round terminates when the step limit is reached, or the *Finish* action is taken by the agent. All predictions are returned, and the complete reasoning process is stored in the action memory.

In our implementation, we use GPT-4o (version 2024-10-01-preview) as the base LLM (Achiam et al., 2023). We enabled the structured output feature to ensure that the model follows the instructions. We train the regression model using the scikit-learn package (Pedregosa et al., 2011) and use packages gget and blitzgsea for data enrichment analysis (Lachmann et al., 2022; Luebbert & Pachter, 2023).

### 3.3 Action Execution

At each step, the agent first selects an action in the pool, as previously described. If the actions are from the *Reasoning* category (Section A.2.1), the agent predicts the results directly. For other action categories, the agent collects and formats the raw execution results. To optimize token usage, we spawn a new agent for the enrichment database selection. This new agent is initialized with the target phenotype information and a list of available database names.

## 4 Experiment Settings

### 4.1 Datasets

We (the authors) preprocessed the following data, and the agent conducted the iterative experiment planning using the given preprocessed data. We selected 11 author-defined phenotypes from a genome-scale Perturb-Seq screen, with 9,867 perturbed genes previously performed in K562 cells (Replogle et al., 2022). The authors used as phenotypes predefined functional gene sets from the CORUM and STRING databases (Giurgiu et al., 2019; Szklarczyk et al., 2021). We included all perturbations as candidates in our experiments to mimic real-world conditions, where the ratio of phenotype-specific hits to the total number of possible perturbations is low. Specifically, of the 9,867 genes perturbed in this screen, the number of hit genes, as defined by the authors (Replogle et al., 2022), across the 11 phenotypes, ranges from 34 to 95. The agent’s target is identifying hit genes for a single phenotype, not considering multiple phenotypes at the same time. Each perturbation is associated with a set of single cell profiles for the individual cells in which the gene is perturbed. Following standard analysis (Replogle et al., 2022), we quantified the impact of each perturbation on phenotypes *S*_*p*_, by applying a z-score transformation to the expression data and clustering perturbations based on the similarity of their associated expression profiles.

Each phenotype is associated with a predefined set of ADGs. The phenotype score for each perturbation was calculated by us as the mean z-score of its ADGs. The dataset’s authors defined the hit genes for a specific phenotype as those whose absolute z-scores were significantly larger than those of other perturbed genes. We used the hit genes provided by the authors as the ground truth in our analysis (Table 1).

**Table 1:**
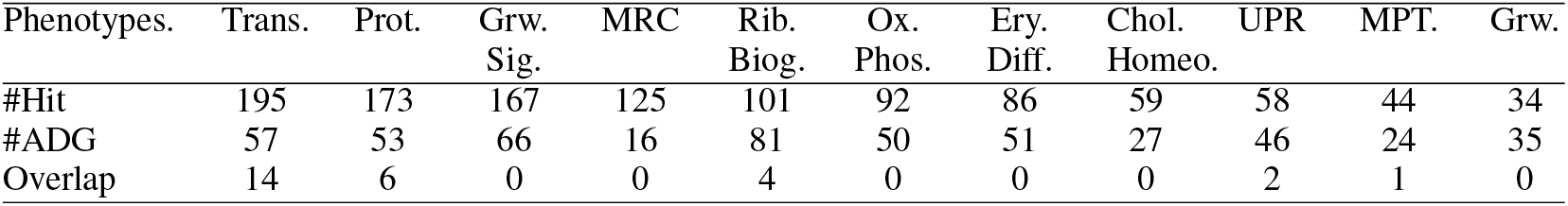
11 Phenotypes in a K562 cell line. Hit genes: genes with impactful effect on each phenotype upon perturbation (columns). ADG: genes whose co-expression constitutes the phenotype, provided as input to PerTurboAgent. Overlap: number of shared hit genes and ADGs, which could thus be leaked to PerTurboAgent. Full phenotypes are in Table 6.

| Phenotypes. | Trans. | Prot. | Grw.<br>Sig. | MRC | Rib.<br>Biog. | Ox.<br>Phos. | Ery.<br>Diff. | Chol.<br>Homeo. | UPR | MPT. | Grw. |
|---|---|---|---|---|---|---|---|---|---|---|---|
| #Hit | 195 | 173 | 167 | 125 | 101 | 92 | 86 | 59 | 58 | 44 | 34 |
| #ADG | 57 | 53 | 66 | 16 | 81 | 50 | 51 | 27 | 46 | 24 | 35 |
| Overlap | 14 | 6 | 0 | 0 | 4 | 0 | 0 | 0 | 2 | 1 | 0 |

### 4.2 Metrics

We defined our primary evaluation metric as the cumulative number of hits *h* identified by the final *N* round, and track the normalized hit ratio, *h*_*ratio*_, where the average *h* is weighted by the number of ground true hits, such that phenotypes with different numbers of total hits can be compared.

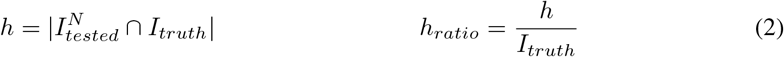

We further define the area under the hit curve (AUC) to assess how well the model finds the overall number of hit perturbations, as:

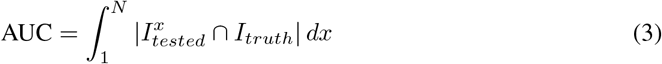

A normalized AUC is defined as the AUC divided by the area under the best hit curve.

### 4.3 Baselines

We compared PerTurboAgent with five methods: *BiodiscoveryAgent, Random, Random+Enrichment, Coreset, and AdversarialBIM* (Kurakin et al., 2018; *Sener & Savarese, 2017)*.

*For BiodiscoveryAgent*, we retained its original strategy but added the ADGs to the task initialization for a fair comparison. We set the number of tries to 20 per round, matching our maximum number of steps. The critique and data analysis modules were enabled, while the literature review module was disabled to prevent errors from queuing irrelevant literature. (Enabling the module led to worse performance.)

The *Random* method randomly selects perturbations from the untested set in each round, whereas the *Random+Enrichment* method includes an enrichment analysis step: if any hit perturbation is identified through random selection, it uses enrichment tools to prioritize the most frequently occurring untested perturbations for the next round. If the enriched set is smaller than needed, random selections fill the gap.

*Coreset* and *AdversarialBIM* are active learning strategies adapted from GeneDisco (Mehrjou et al., 2021). These methods train Bayesian Neural Networks (BNNs) on existing data and use Bayesian optimization to rank and select untested perturbations in each round. We conducted five replicates per method for each task.

## 5 Results

In this section, we first present benchmarking results for all methods and then analyze PerTurboAgent’s behavior to improve interpretability, followed by ablation studies that highlight the importance of different agent actions. Finally, we assess our framework’s ability to generalize using both closed-source and open-source models.

### 5.1 Benchmarking

PerTurboAgent outperformed all baseline methods in terms of both mean number of hits and hit ratio (Table 2). *Random+Enrichment* served as a strong baseline, demonstrating the benefits of simple data analysis with prior knowledge in guiding experimental design. While *BioDiscoveryAgent* uses similar data analysis, its reliance on fixed queries to the Reactome database and occasional query instability may have hindered its performance. PerTurboAgent excelled in the top 6 phenotype tasks with more hits (Figure 2), such as “Translation,” “Proteasome,” and “Mitochondrial Respiratory Chain.” For the five phenotype tasks with fewer hits, PerTurboAgent still surpassed the second-best methods (hit ratio: 0.27 vs. 0.23), although hit curves for all LLM-based methods showed greater variance (Figure 5). This may result from fewer early-round hits, reducing the stabilizing effect of feedback-driven refinement in subsequent rounds. PerTurboAgent also showed a strong performance gain on AUC metrics, reflecting its efficiency in quickly identifying hits.

**Table 2:** Performance benchmarks. Each entry is the average across all phenotypes.

|  | PerTurboAgent | BioDisAgent | Enrich+Rand | Coreset | AdvBIM | Random |
|---|---|---|---|---|---|---|
| Hit number | **51.127** | 24.164 | 30.727 | 5.709 | 7.273 | 6.091 |
| Hit ratio | **0.440** | 0.240 | 0.255 | 0.055 | 0.066 | 0.057 |
| AUC | **290.236** | 131.455 | 171.873 | 29.864 | 36.545 | 28.545 |
| Normed AUC | **0.280** | 0.154 | 0.158 | 0.033 | 0.037 | 0.030 |

**Figure 2:**
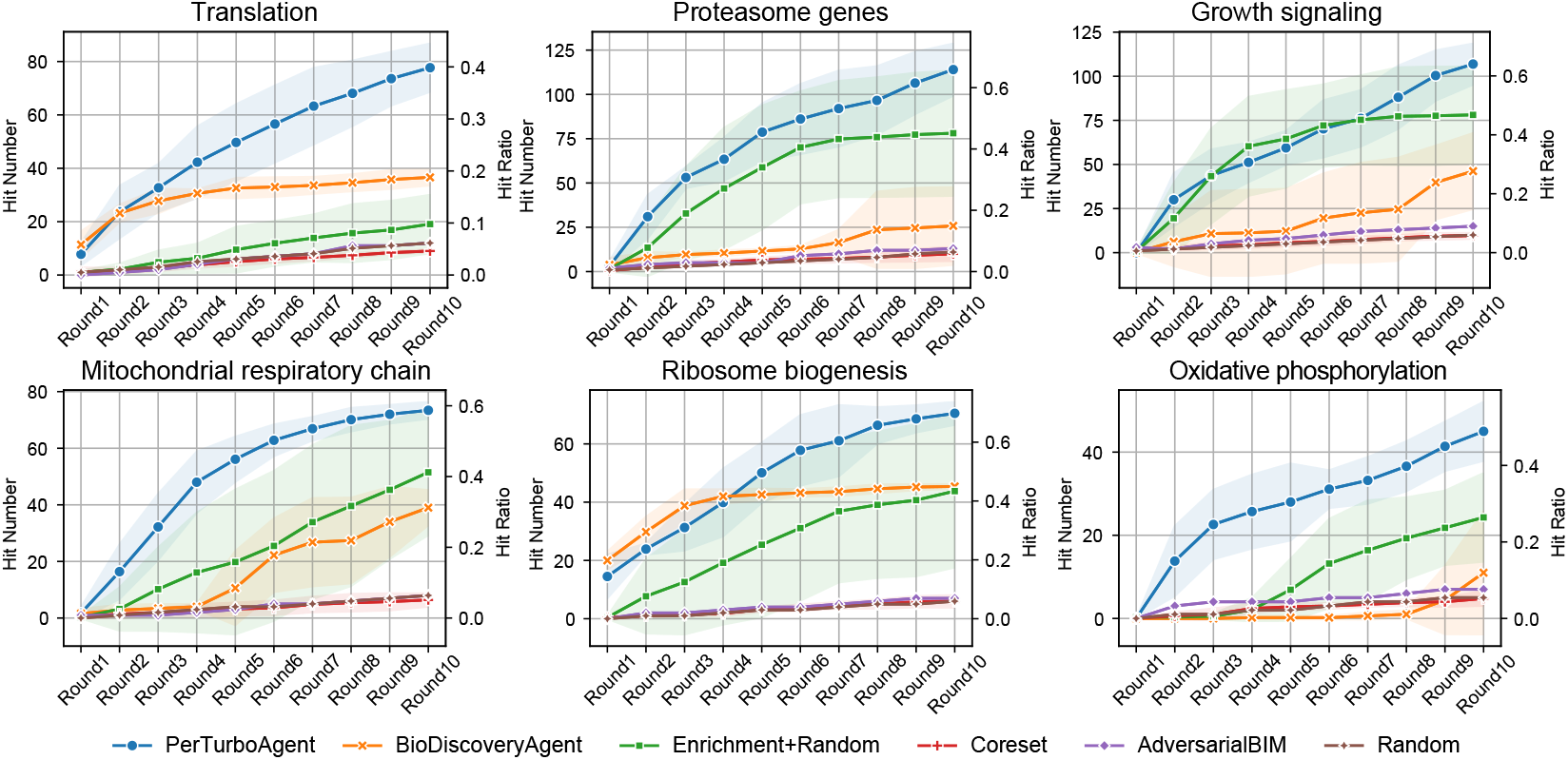
Accrual of hits over rounds. Average number of hits (left y-axis) and hit ratio (right y-axis) in each round (x-axis) from 5 runs for six phenotypes (panels) with high hit numbers. Error bars: standard deviation. Other phenotypes are in Fig. 5

To determine if ADGs overly influenced the agent’s predictions in the task initialization prompt, we tracked the overlap between cumulative predictions and the ADG set across all rounds (Table 7). Initially, the overlap increased rapidly and then stabilized. The agent predicted a large fraction of the ADGs for the translation (47 of 57) and cholesterol homeostasis (22 of 27) tasks. While in the translation task, there was a high overlap between ground-truth hits and ADGs, in other tasks, fewer than half of the ADGs were predicted. These observations suggest that PerTurboAgent can leverage multiple information sources rather than solely relying on the prompt.

### 5.2 Behavior Analysis

Behavioral analysis of PerTurboAgent revealed that internal knowledge–related actions were the most frequently executed (Figure 3a), and that some action trends shifted over the rounds (Figure 3b). In particular, the agent increasingly used the machine learning (ML) prediction model in later rounds, suggesting that it recognizes the value of additional training data for enhancing the ML model’s predictive performance. In contrast, the frequency of GSEA on control cells declined over rounds, aligning with the intuitive notion that information derived from hits is more valuable than that from reference controls. Most other actions maintained a stable frequency across rounds.

**Figure 3:**
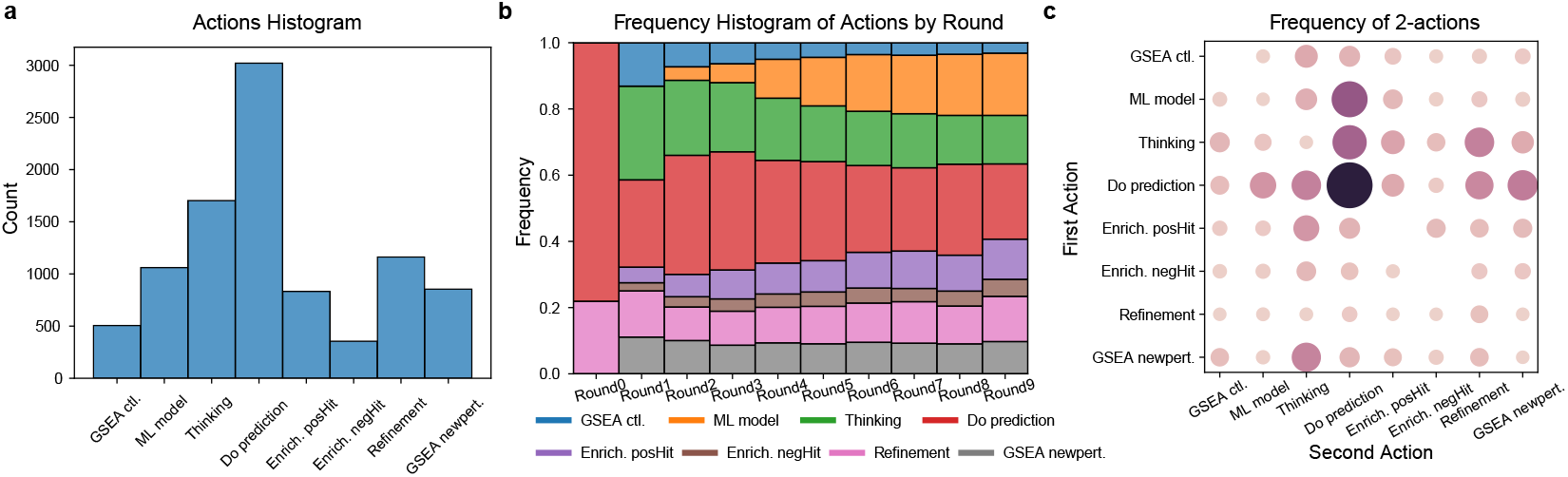
Behavior analysis of PerTurboAgent. **a** Number of actions (y axis) in each category (x axis). **b** Proportion of actions (y-axis)from each category (color) in each round (x-axis). **c** Consecutive action pairs. Frequency (dot color of size) of the next action (columns) following action from each category (rows).

Examining the frequency of consecutive action pairs (Figure 3c) revealed some action dependencies, including the expected pattern of consecutive prediction steps, and the combination of “GSEA on new perturbations” followed by “Reflecting on results stored in memory”, showing that when confronted with complex, data-rich outputs such as GSEA results, the agent engages in deeper reasoning, carefully integrating and interpreting the information before proceeding. For example, a portion of the Think Log for the protein translation task (Figure 4) illustrates the agent’s structured reflection on previous results, their connection to GSEA findings, and the subsequent planning of future actions or predictions.

**Figure 4:**
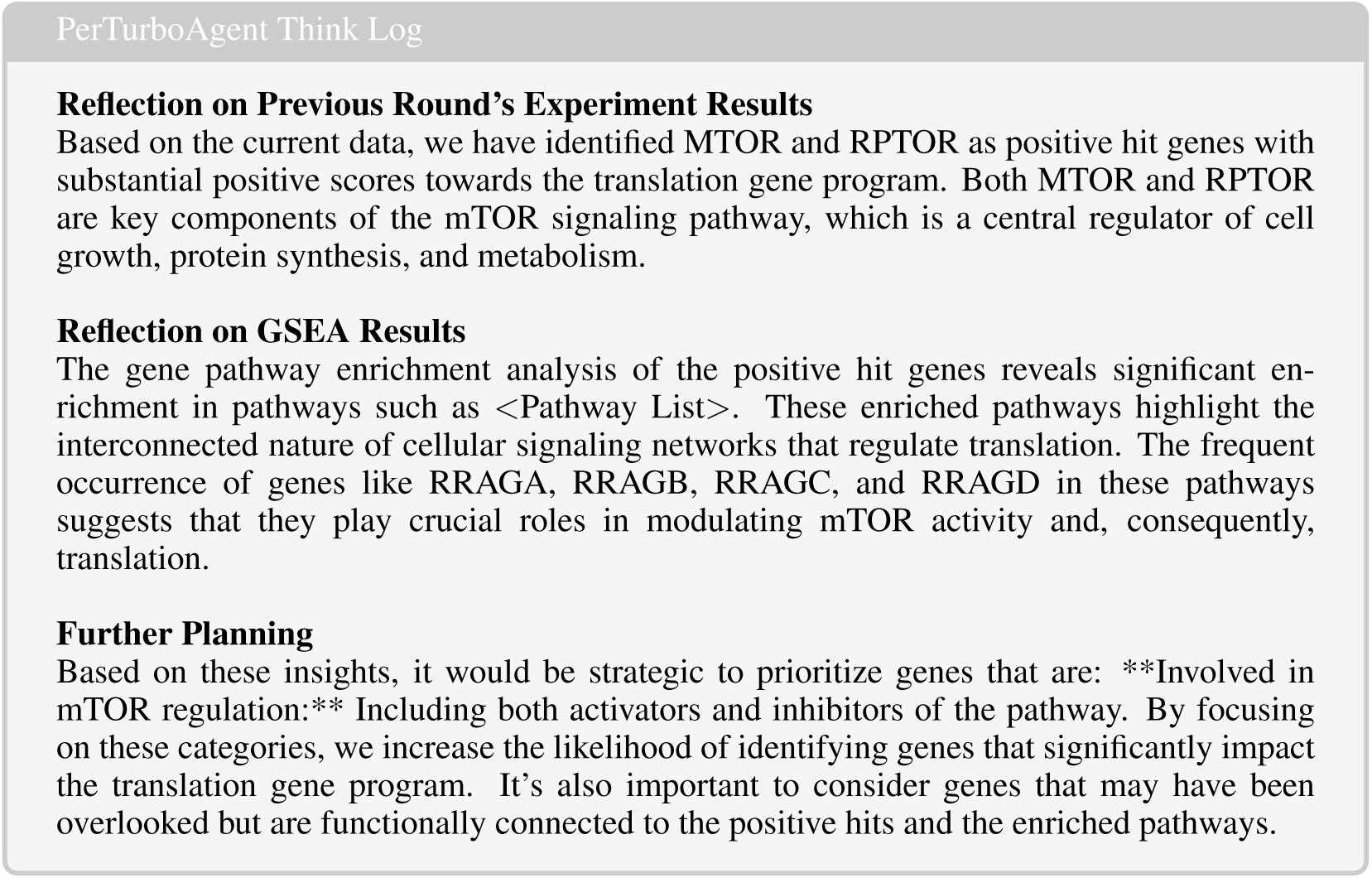
Example log of consecutive PerTurboAgent actions. Reflection after GSEA by PerTurboAgent on the “Translation” phenotype.

#### 5.3 Ablation Study

To assess the impact of different action classes on the Agent’s performance, we first considered the impact of removing its self-reflection action, which, alongside reflective reasoning, is part of the prediction action (A.2.2). The removal led to a decline in all performance metrics, indicating that explicit reasoning enhances both interpretability and prediction performance, even with a Chain-of-Thought (CoT) format (Table 3).

**Table 3:** Action ablation study.

|  | Baseline | w/o<br>Thinking | w/o<br>HitGSEA | w/o<br>CtrlGSEA | w/o<br>GSEA | w/o<br>Enrich | w/o<br>Enrich&GSEA | w/o<br>ML |
|---|---|---|---|---|---|---|---|---|
| Hit number | **51.127** | 48.473 | 46.145 | 49.523 | 45.891 | 46.909 | 41.682 | 50.555 |
| Hit ratio | **0.440** | 0.411 | 0.422 | 0.408 | 0.391 | 0.402 | 0.371 | 0.433 |
| AUC | **290.236** | 265.318 | 281.500 | 231.709 | 238.073 | 278.445 | 191.034 | 284.527 |
| Normed AUC | **0.280** | 0.259 | 0.260 | 0.229 | 0.251 | 0.264 | 0.194 | 0.280 |

We also explored the effects of different data analysis actions. Removing the GSEA action on new hits and control cell lines reduced performance, with the GSEA on new hits having a more substantial impact. Eliminating all GSEA actions further degraded results. A similar decline occurred when we removed all enrichment steps, with the largest performance drop resulting from removing enrichment actions on gene sets and expression profiles. Removing the training perturbation prediction model action caused only a slight performance decrease. Despite small performance changes compared to other actions, this small reduction suggests the agent still gleaned useful information from the model’s predictions.

We further evaluated the impact of introducing new actions. Both removing the original constraint on the ML model to allow training to begin as early as the second round and testing an enrichment action on non-hit perturbations similarly harmed performance (Table 4). We hypothesize that because our task involves fewer than 200 hits, enriching non-relevant perturbations likely introduces noise, reducing the signal-to-noise ratio.

**Table 4:** Ablation study of adding more actions.

|  | Baseline | all ML | w/ EnrichNonHit |
|---|---|---|---|
| Hit number | **51.127** | 48.409 | 49.018 |
| Hit ratio | **0.440** | 0.412 | 0.403 |
| AUC | **290.236** | 241.420 | 273.073 |
| Normed AUC | **0.280** | 0.239 | 0.256 |

### 5.4 Performance of Different LLMs

**Table 5:** Performance of different base LLMs.

|  | GPT-4o | GPT-4 | Claude35Sonnet | o1-mini | o1-preview | Qwen2.5-72B |
|---|---|---|---|---|---|---|
| Hit number | 51.127 | 47.364 | 51.500 | 34.409 | **57.273** | 48.364 |
| Hit ratio | 0.440 | 0.440 | 0.441 | 0.351 | **0.521** | 0.466 |
| AUC | 290.236 | 255.364 | 279.023 | 190.727 | **311.409** | 246.955 |
| Normed AUC | 0.280 | 0.271 | 0.278 | 0.231 | **0.330** | 0.265 |
| \$ / 1M Token | \$10.00 | \$60.00 | \$15.00 | \$12.00 | \$60.00 | \$0.40 |

Finally, we evaluated the impact of various base LLMs on PerTurboAgent’s performance, testing both closed-source models (GPT-4o, GPT-4, Claude 3.5 Sonnet 2024-10-22, Claude 3.5 Haiku, o1-mini, o1-preview) and open-source models (Qwen2.5-72B, Llama 3.3-70B) (Yang et al., 2024; Dubey et al., 2024), deployed via the vLLM (Kwon et al., 2023). Overall, our framework was compatible with various underlying models. The o1-preview model delivered the strongest performance but incurred the highest cost. Rapid-response models like Claude-Haiku and o1-mini did not achieve strong results. The Qwen2.5-72B model performed similarly to GPT-4, while Llama 3.3-70B failed to follow instructions and consequently produced no usable outcomes.

## 6 Discussion

Sequential perturb-seq experiments can more efficiently identify perturbations linked to a target phenotype for a fixed overall experimental footprint. A primary challenge is selecting perturbations that integrate results from previous rounds with existing domain knowledge. Traditional methods often struggle with representing prior knowledge numerically. While LLMs show promise for sequential experiment design, the role of traditional methods and deeper data analyses, like gene expression profiles, in enhancing LLM-driven agents remains under-explored.

PerTurboAgent addresses this challenge by using an action memory and a diverse pool of specialized actions. The agent autonomously plans a multi-step process, integrating conventional data analyses, machine learning models, and its own reasoning. This iterative approach provides interpretable action trajectories and actionable insights, significantly improving perturb-seq experiment design.

Future work on PerTurboAgent could include integrating advanced perturbation prediction models, such as GEARS, or fine-tuning the pre-trained foundation models (e.g., scGPT, scFoundation) to improve phenotype effect estimation (Roohani et al., 2024a; Hao et al., 2024; Cui et al., 2024). In this study, although PerTurboAgent was able to conduct autonomous analysis and planning for perturb-seq data, it still required manual curation and preprocessing, and the target phenotypes we provided were for well-defined biological processes. Extending capabilities to *ab initio* phenotype discovery would enable autonomous identification of significant phenotypes without predefined targets. Incorporating multiple agents for action evaluation and feedback could enhance efficiency, as suggested by recent work (Yu et al., 2024). Additionally, fine-tuning open-source models using reinforcement learning on past experimental data shows promise for further performance gains (Ouyang et al., 2022; Rafailov et al., 2023), offering an alternative to proprietary models.

## Acknowledgments

This work was done while M.H. was an intern at Genentech. Y.L., H.W., G.S., and A.R. are employees of Genentech, a member of the Roche Group. A.R. has equity in Roche. A.R. is a co-founder and equity holder of Celsius Therapeutics, an equity holder in Immunitas, and until 31 July 2020 was on the scientific advisory board of ThermoFisher Scientific, Syros Pharmaceuticals, Neogene Therapeutics and Asimov.

## A Appendix

### A.1 Initialization Prompt

In this section, we present the prompt we used to help the LLMs understand the tasks and actions, as well as the specific prompt formats tailored for each action. To further control the output and ensure accurate parsing, we utilized the structured output feature of the OpenAI API. For other models, we incorporated XML-based grammar directly into the prompt to improve the reliability of text parsing. All italic text in the prompt represents variables.

##### Task initialization Prompt

You are a scientist working on problems in drug discovery. Research Problem: I’m planning to run a CRISPR screen to identify CRISPERI (knockdown) gene perturbations that affect *phenotypename* related gene programs. There are 9,867 possible genes to perturb and I can only perturb *numgenes* genes at a time. For each perturbation, I’m able to measure out the z score of the target gene program in perturbed data. The z score value of gene program is the mean z score value of these genes: *genelist*. For each gene, we calculated its z score by using the mean and standard deviation from the reference control data. Your task is predicting and prioritizing genes for the gene panel design in each round to find the most relevant genes (hit genes) to the research problem.

##### Action Loading Prompt

There are several actions you can choose from to learn more about the problem and previous data. You can request the following: 1. *ActionName*1: *Description*1 2. *ActionName*2: *Description*2 … Note that you can choose an option more than once, but each time you can only choose one. In total, you can choose options up to 20 times. All valid actions are *V alidActions*. Provide only your most desired choice in the format *<*STEP*>*Number*<*/STEP*>* with no more than two sentences of explanation.

##### Different strategy prompt

Exploration: *“One strategy you can follow is to focus on trying a very diverse set of genes to get a sense of which types of genes affect the research problem the most*.*”* Exploitation: *“One strategy you can follow is to double down on pathways that include many hits to increase the cumulative hit rate*.*”*

### A.2 Action Pool

#### A.2.1 Group 1

##### Action 1: Predicting new perturbations

*Description*: Indicate that you are ready to output your predictions. You will provide and prioritize genes following a specific format. Call this action when your prediction does not exceed the number of genes you need to predict. If you want to refine your existing predictions, choose action 9.

*Action Prompt*: You have to predict *numgenes* genes. Use HGNC gene naming convention. DO NOT PREDICT GENES THAT HAVE ALREADY BEEN TESTED OR PREDICTED.

Respond in this format exactly: 1. Reflection: Thoughts on previous results and next steps. 2. Research Plan: The full high level research plan, with current status and reasoning behind each proposed approach. It should be at most 5 sentences. 3. Solution: A list of predicted genes: [*<*Gene name 1*>, <*Gene name 2*>*, …]

##### Action 2: Reflecting results stored in memory

*Description*: Reflect on the current data and output your analysis, providing insights for further investigation.

*Action Prompt*: Show your reflections here.

##### Action 3: Refining existing predictions

*Description*: Refine your prediction based on new observations and feedback.

*Action Prompt*: Your current prediction has *totalnum* genes. Think carefully, critique the current plan and propose the genes you want to remove and add. All other genes will be kept. Use HGNC gene naming convention. DO NOT PREDICT GENES THAT HAVE ALREADY BEEN TESTED. Please do not make changes if there is no need to make a change.

Respond in this format exactly: 1. Critique: include all relevant details of the critique. 2. Solution-Removal: Give a list of genes you want to remove from the current prediction separated by commas in this format: [*<*Gene name 1*>*,…] 3. SolutionAddition: Give a list of genes you want to add to the current prediction separated by commas in this format: [*<*Gene name 1*>*, …]

##### Action 4: Finish

*Description*: Finish the current round and output the final prediction. If the length of the current solution exceeds the number of genes you need to predict, the first *numgenes* genes will be selected as the final prediction.

*Action Prompt*: N/A

#### A.2.2 Group 2

##### Action 5: Training pert. prediction model

*Description*: Request the top *numgenes* genes most likely to have a high (high predicted absolute value) impact on the target gene program, predicted by a regression model. The model will be trained on previous experimental data, the input of the model is the embedding of one perturbation, and the output is the predicted score of the target gene expression program under the input perturbation.

Then we trained the model as depicted in the method section. The format of the output result is: The top *numgene//*2 prediction with max score are *posgene*. The top *numgene//*2 prediction with min score are *neggene*

#### A.2.3 Group 3

##### Action 6: Enrichment Analysis

*Description*: Request the gene pathway enrichment analysis on hit genes with positive/negative scores. We will return up to the top 10 enriched pathways with p-values smaller than 0.05, based on different libraries. And we will provide the most frequent valid genes shared in these enriched pathways.

*Database selection prompt*: You are a scientist working on problems in drug discovery. Research Problem: *research*_*p*_*roblem* For now, you are asked to perform the gene pathway enrichment analysis on the control cell. Choose three desired libraries for the analysis. all available libraries are: *GeneSigDB, KEGG*_2_021_*H*_*uman*, … Return your answer in the format:*<*LIB1*>*libname1*<*/LIB1*> <*LIB2*>*libname2*< /*LIB2*> <*LIB3*>*libname3*< /*LIB3*>*.

*Result format*: We performed the enrichment analysis based on three libraries. For the first library *libname*1, the top enriched pathway is *sigpat*1. For the second library *libname*2, the top enriched pathway is *sigpath*2. For the third library *libname*3, the top enriched pathway is *sigpath*3. These pathways span various biological processes and may be linked to different diseases or cellular functions. The gene names provided here have been identified as the candidates for most frequently occurring genes across all enriched pathways: *genelist*

##### Action 7: GSEA Analysis

*Description*: Request the gene set enrichment analysis on hit genes that have been experimented on in the previous round. We will return up to the top 3 enriched pathways with p-values smaller than 0.05 for each hit perturbation.

*Database selection prompt*: Same as the Action 6

*Result format*: We performed GSEA based on three libraries. (same as the Action 6)

### A.3 Supplementary Results

**Table 6:** Full list of shortened phenotype name.

| **Abbreviation** | **Phenotype Name** |
|---|---|
| Trans. | Translation; |
| Prot. | Proteasome Genes; |
| Grw. Sig. | Growth Signaling; |
| MRC | Mitochondrial Respiratory Chain; |
| Rib. Biog. | Ribosome Biogenesis; |
| Ox. Phos. | Oxidative Phosphorylation; |
| Ery. Diff. | Erythroid Differentiation; |
| Chol. Homeo. | Cholesterol Homeostasis; |
| UPR | Unfolded Protein Response/mTORC1 Signaling; |
| MPT. | Mitochondrial Protein Translocation and Oxidative Phosphorylation; |
| Grw. | Growth/Targets of Myc. |

**Figure 5:**
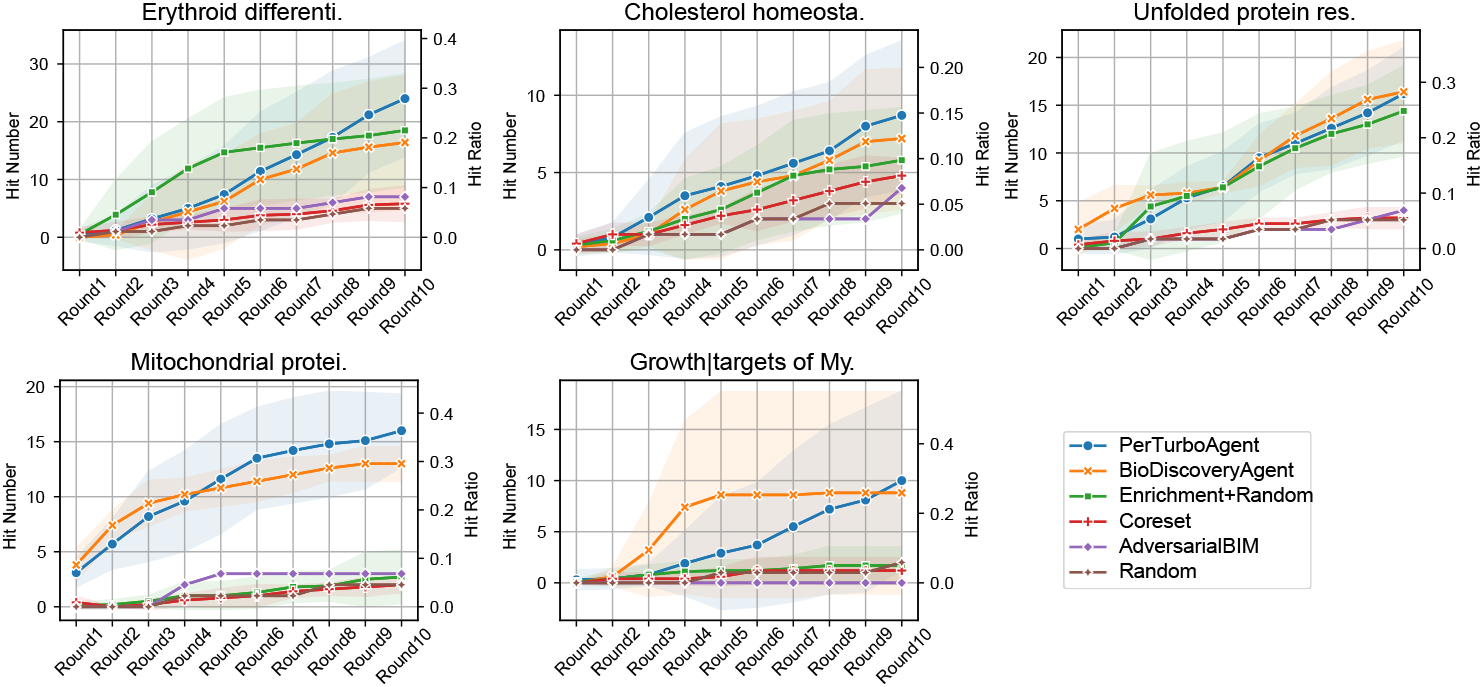
Hit curve of the rest five phenotype tasks

**Table 7:** Overlap between hit genes and phenotype associated descriptive genes.

| **Rounds** | *Trans.* | *Prot.* | *Grw. Sig.* | *MRC* | *Rib. Biog.* | *Ox. Phos.* |
|---|---|---|---|---|---|---|
| **0** | 1.1 | 0.7 | 13.8 | 0.0 | 6.6 | 6.0 |
| **1** | 29.3 | 8.0 | 14.1 | 0.0 | 9.0 | 6.3 |
| **2** | 39.3 | 10.9 | 14.1 | 0.0 | 16.8 | 7.4 |
| **3** | 41.7 | 13.4 | 14.1 | 0.0 | 21.0 | 9.0 |
| **4** | 43.7 | 17.6 | 14.2 | 0.0 | 24.3 | 9.9 |
| **5** | 45.1 | 19.5 | 14.6 | 0.2 | 25.8 | 10.4 |
| **6** | 45.5 | 20.6 | 14.7 | 0.4 | 27.5 | 11.2 |
| **7** | 46.0 | 21.8 | 15.1 | 0.6 | 29.7 | 12.7 |
| **8** | 46.3 | 22.9 | 15.2 | 0.6 | 31.2 | 13.9 |
| **9** | 46.6 | 24.4 | 15.7 | 0.6 | 32.5 | 14.7 |
| **#ADG** | 57 | 53 | 66 | 16 | 81 | 50 |
| **#Overlap** | 14 | 6 | 0 | 0 | 4 | 0 |

**Table 8:** Hit ratio performance across all methods.

|  | *Trans.* | *Prot.* | *Grw. Sig.* | *MRC* | *Rib. Biog.* | *Ox. Phos.* | *Ery. Diff.* | *Chol. Homeo.* | *UPR* | *MPT.* | *Grw.* | *Mean* |
|---|---|---|---|---|---|---|---|---|---|---|---|---|
| **PerTurboAgent** | **0.398** | **0.659** | **0.640** | **0.587** | **0.698** | **0.489** | **0.279** | **0.147** | 0.279 | **0.364** | **0.294** | **0.440** |
| **BioDisAgent** | 0.188 | 0.149 | 0.277 | 0.312 | 0.450 | 0.120 | 0.191 | 0.122 | **0.283** | 0.295 | 0.259 | 0.240 |
| **Enrich+Rand** | 0.098 | 0.451 | 0.468 | 0.412 | 0.434 | 0.264 | 0.215 | 0.098 | 0.248 | 0.061 | 0.050 | 0.255 |
| **Coreset** | 0.046 | 0.058 | 0.057 | 0.051 | 0.059 | 0.052 | 0.067 | 0.081 | 0.055 | 0.045 | 0.035 | 0.055 |
| **AdvBIM** | 0.062 | 0.075 | 0.090 | 0.064 | 0.069 | 0.076 | 0.081 | 0.068 | 0.069 | 0.068 | 0.000 | 0.066 |
| **Random** | 0.062 | 0.064 | 0.060 | 0.064 | 0.059 | 0.054 | 0.058 | 0.051 | 0.052 | 0.045 | 0.059 | 0.057 |

**Table 9:**
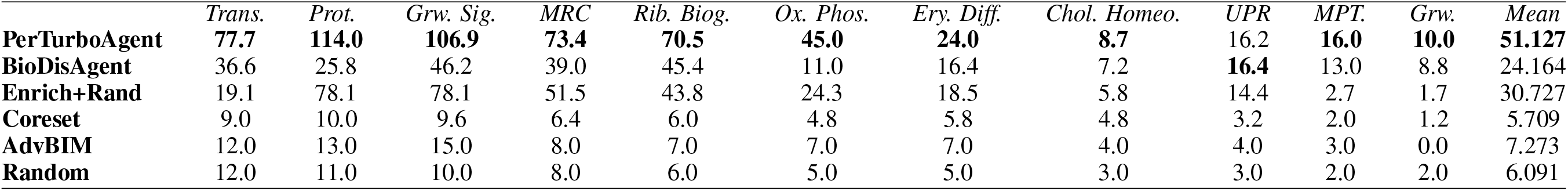
Hit number performance across all methods.

**Table 10:** Normed area under hit curve (AUC) scores across all methods.

|  | *Trans.* | *Prot.* | *Grw. Sig.* | *MRC* | *Rib. Biog.* | *Ox. Phos.* | *Ery. Diff.* | *Chol. Homeo.* | *UPR* | *MPT.* | *Grw.* | *Mean* |
|---|---|---|---|---|---|---|---|---|---|---|---|---|
| **PerTurboAgent** | **0.280** | **0.457** | **0.405** | **0.422** | **0.496** | **0.313** | **0.122** | **0.075** | **0.138** | **0.258** | 0.117 | **0.280** |
| **BioDisAgent** | 0.170 | 0.090 | 0.120 | 0.138 | 0.407 | 0.015 | 0.097 | 0.063 | 0.156 | 0.240 | **0.193** | 0.154 |
| **Enrich+Rand** | 0.056 | 0.336 | 0.375 | 0.201 | 0.264 | 0.114 | 0.150 | 0.054 | 0.131 | 0.029 | 0.034 | 0.158 |
| **Coreset** | 0.029 | 0.038 | 0.038 | 0.030 | 0.033 | 0.029 | 0.040 | 0.042 | 0.036 | 0.022 | 0.024 | 0.033 |
| **AdvBIM** | 0.035 | 0.048 | 0.057 | 0.032 | 0.041 | 0.051 | 0.050 | 0.024 | 0.027 | 0.047 | 0.000 | 0.037 |
| **Random** | 0.036 | 0.035 | 0.035 | 0.033 | 0.030 | 0.030 | 0.031 | 0.027 | 0.028 | 0.023 | 0.020 | 0.030 |

**Table 11:** Absolute area under hit curve (AUC) scores across all methods.

|  | *Trans.* | *Prot.* | *Grw. Sig.* | *MRC* | *Rib. Biog.* | *Ox. Phos.* | *Ery. Diff.* | *Chol. Homeo.* | *UPR* | *MPT.* | *Grw.* | *Mean* |
|---|---|---|---|---|---|---|---|---|---|---|---|---|
| **PerTurboAgent** | **452.750** | **665.850** | **572.550** | **462.050** | **441.600** | **254.900** | 93.200 | **39.800** | 72.000 | **102.250** | 35.650 | **290.236** |
| **BioDisAgent** | 275.200 | 131.600 | 170.200 | 151.500 | 362.500 | 12.100 | 73.800 | 33.500 | **81.400** | 95.200 | **59.000** | 131.455 |
| **Enrich+Rand** | 91.250 | 489.850 | 529.400 | 219.500 | 234.800 | 92.800 | **114.200** | 28.550 | 68.250 | 11.600 | 10.400 | 171.873 |
| **Coreset** | 46.500 | 55.900 | 53.500 | 32.300 | 29.800 | 23.400 | 30.300 | 22.400 | 18.600 | 8.600 | 7.200 | 29.864 |
| **AdvBIM** | 56.000 | 69.500 | 80.000 | 34.500 | 36.500 | 41.500 | 38.500 | 13.000 | 14.000 | 18.500 | 0.000 | 36.545 |
| **Random** | 58.500 | 51.000 | 49.500 | 36.000 | 27.000 | 24.500 | 23.500 | 14.500 | 14.500 | 9.000 | 6.000 | 28.545 |

**Table 12:** Hit number performance on ablation study.

| **Methods** | *Trans.* | *Prot.* | *Grw. Sig.* | *MRC* | *Rib. Biog.* | *Ox. Phos.* | *Ery. Diff.* | *Chol. Homeo.* | *UPR/mTORC1* | *MPT.* | *Grw./Myc* | *Mean* |
|---|---|---|---|---|---|---|---|---|---|---|---|---|
| PerTurboAgent | 77.7 | 114.0 | 106.9 | 73.4 | 70.5 | 45.0 | 24.0 | 8.7 | 16.2 | 16.0 | 10.0 | 51.127 |
| w/ EnrichNonHit | 74.0 | 103.8 | 112.8 | 73.0 | 68.8 | 44.0 | 22.4 | 6.6 | 19.2 | 14.0 | 0.6 | 49.018 |
| all ML | 67.0 | 108.0 | 110.3 | 65.5 | 70.3 | 38.3 | 29.8 | 6.5 | 13.0 | 17.0 | 7.0 | 48.409 |
| w/o ML | 82.0 | 99.3 | 111.5 | 72.5 | 72.0 | 44.4 | 21.6 | 10.0 | 19.0 | 18.8 | 5.0 | 50.555 |
| w/o NewHitGSEA | 67.0 | 88.4 | 99.2 | 66.6 | 59.0 | 38.4 | 28.6 | 7.8 | 19.8 | 17.2 | 15.6 | 46.145 |
| w/o CtrlGSEA | 67.8 | 112.3 | 112.8 | 73.8 | 69.8 | 42.5 | 26.0 | 9.8 | 17.5 | 9.8 | 3.0 | 49.523 |
| w/o GSEA | 66.2 | 94.2 | 100.0 | 72.6 | 65.4 | 37.0 | 24.8 | 7.0 | 16.2 | 18.8 | 2.6 | 45.891 |
| w/o Enrich | 66.8 | 108.2 | 95.2 | 66.4 | 65.6 | 45.0 | 23.0 | 8.8 | 18.2 | 10.6 | 8.2 | 46.909 |
| w/o GSEA&Enrich | 61.5 | 91.3 | 82.3 | 49.8 | 59.3 | 40.8 | 24.5 | 6.8 | 18.8 | 15.3 | 8.5 | 41.682 |
| w/o Thinking | 74.2 | 107.6 | 103.2 | 68.2 | 69.8 | 40.8 | 22.6 | 8.2 | 17.2 | 15.6 | 5.8 | 48.473 |

**Table 13:**
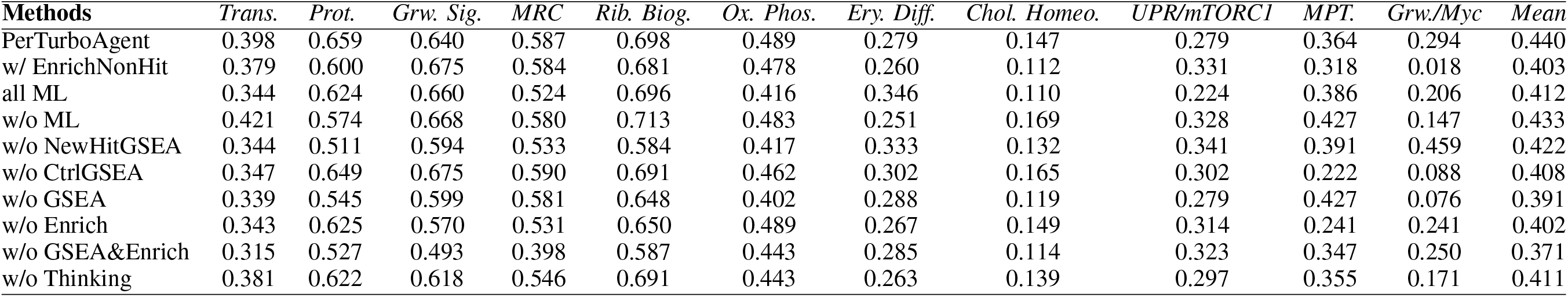
Hit ratio performance on ablation study.

| **Methods** | *Trans.* | *Prot.* | *Grw. Sig.* | *MRC* | *Rib. Biog.* | *Ox. Phos.* | *Ery. Diff.* | *Chol. Homeo.* | *UPR/mTORC1* | *MPT.* | *Grw./Myc* | *Mean* |
|---|---|---|---|---|---|---|---|---|---|---|---|---|
| PerTurboAgent | 0.398 | 0.659 | 0.640 | 0.587 | 0.698 | 0.489 | 0.279 | 0.147 | 0.279 | 0.364 | 0.294 | 0.440 |
| w/ EnrichNonHit | 0.379 | 0.600 | 0.675 | 0.584 | 0.681 | 0.478 | 0.260 | 0.112 | 0.331 | 0.318 | 0.018 | 0.403 |
| all ML | 0.344 | 0.624 | 0.660 | 0.524 | 0.696 | 0.416 | 0.346 | 0.110 | 0.224 | 0.386 | 0.206 | 0.412 |
| w/o ML | 0.421 | 0.574 | 0.668 | 0.580 | 0.713 | 0.483 | 0.251 | 0.169 | 0.328 | 0.427 | 0.147 | 0.433 |
| w/o NewHitGSEA | 0.344 | 0.511 | 0.594 | 0.533 | 0.584 | 0.417 | 0.333 | 0.132 | 0.341 | 0.391 | 0.459 | 0.422 |
| w/o CtrlGSEA | 0.347 | 0.649 | 0.675 | 0.590 | 0.691 | 0.462 | 0.302 | 0.165 | 0.302 | 0.222 | 0.088 | 0.408 |
| w/o GSEA | 0.339 | 0.545 | 0.599 | 0.581 | 0.648 | 0.402 | 0.288 | 0.119 | 0.279 | 0.427 | 0.076 | 0.391 |
| w/o Enrich | 0.343 | 0.625 | 0.570 | 0.531 | 0.650 | 0.489 | 0.267 | 0.149 | 0.314 | 0.241 | 0.241 | 0.402 |
| w/o GSEA&Enrich | 0.315 | 0.527 | 0.493 | 0.398 | 0.587 | 0.443 | 0.285 | 0.114 | 0.323 | 0.347 | 0.250 | 0.371 |
| w/o Thinking | 0.381 | 0.622 | 0.618 | 0.546 | 0.691 | 0.443 | 0.263 | 0.139 | 0.297 | 0.355 | 0.171 | 0.411 |

**Table 14:**
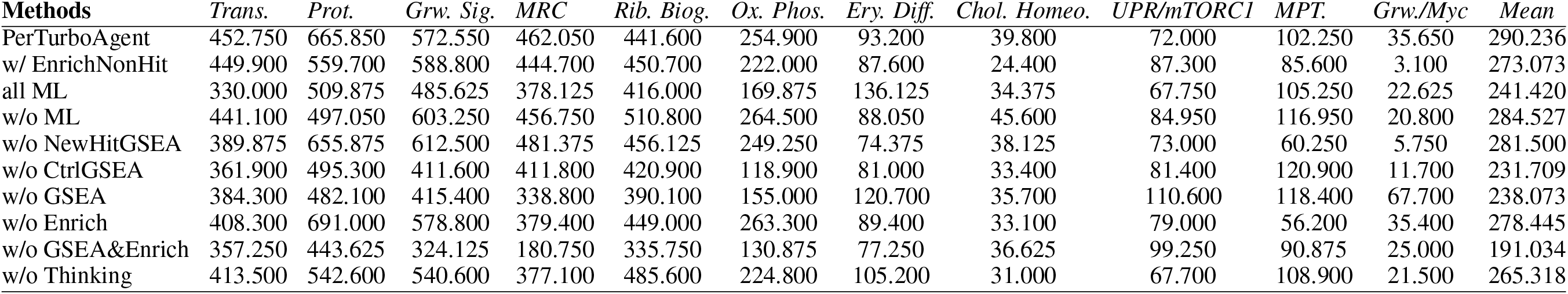
Area under hit curve (AUC) performance on ablation study.

**Table 15:** Normed area under hit curve (AUC) performance on ablation study.

| **Methods** | *Trans.* | *Prot.* | *Grw. Sig.* | *MRC* | *Rib. Biog.* | *Ox. Phos.* | *Ery. Diff.* | *Chol. Homeo.* | *UPR/mTORC1* | *MPT.* | *Grw./Myc* | *Mean* |
|---|---|---|---|---|---|---|---|---|---|---|---|---|
| PerTurboAgent | 0.280 | 0.457 | 0.405 | 0.422 | 0.496 | 0.313 | 0.122 | 0.075 | 0.138 | 0.258 | 0.117 | 0.280 |
| w/ EnrichNonHit | 0.278 | 0.384 | 0.417 | 0.406 | 0.506 | 0.273 | 0.115 | 0.046 | 0.167 | 0.216 | 0.010 | 0.256 |
| w/o ML | 0.272 | 0.341 | 0.427 | 0.417 | 0.574 | 0.325 | 0.115 | 0.086 | 0.163 | 0.295 | 0.068 | 0.280 |
| all ML | 0.204 | 0.350 | 0.344 | 0.345 | 0.467 | 0.209 | 0.178 | 0.065 | 0.130 | 0.266 | 0.074 | 0.239 |
| w/o NewHitGSEA | 0.241 | 0.450 | 0.434 | 0.440 | 0.512 | 0.306 | 0.097 | 0.072 | 0.140 | 0.152 | 0.019 | 0.260 |
| w/o CtrlGSEA | 0.223 | 0.340 | 0.291 | 0.376 | 0.473 | 0.146 | 0.106 | 0.063 | 0.156 | 0.305 | 0.038 | 0.229 |
| w/o GSEA | 0.237 | 0.331 | 0.294 | 0.310 | 0.438 | 0.190 | 0.158 | 0.067 | 0.212 | 0.299 | 0.221 | 0.251 |
| w/o Enrich | 0.252 | 0.474 | 0.410 | 0.347 | 0.504 | 0.323 | 0.117 | 0.062 | 0.151 | 0.142 | 0.116 | 0.264 |
| w/o GSEA&Enrich | 0.221 | 0.304 | 0.229 | 0.165 | 0.377 | 0.161 | 0.101 | 0.069 | 0.190 | 0.229 | 0.082 | 0.194 |
| w/o Thinking | 0.255 | 0.372 | 0.383 | 0.345 | 0.545 | 0.276 | 0.138 | 0.058 | 0.130 | 0.275 | 0.070 | 0.259 |

### A.4 Full log of single round

##### Task initialization Prompt

You are a scientist working on problems in drug discovery. Research Problem: I’m planning to run a CRISPR screen to identify CRISPERI (knockdown) gene perturbations that affect translation related gene programs. There are 9,867 possible genes to perturb and I can only perturb 64 genes at a time. For each perturbation, I’m able to measure out the z score of the target gene program in perturbed data. The z score value of gene program is the mean z score value of these genes: [‘ACTG1’, ‘CCNG1’, ‘COX7C’, ‘EEF2’, ‘EIF2S3’, ‘EIF3F’, ‘EIF4B’, ‘HIGD2A’, ‘HNRNPA1’, ‘POLR1D’, ‘RPL10’, ‘RPL10A’, ‘RPL12’, ‘RPL13’, ‘RPL13A’, ‘RPL14’, ‘RPL15’, ‘RPL18’, ‘RPL18A’, ‘RPL19’, ‘RPL22’, ‘RPL23’, ‘RPL23A’, ‘RPL26’, ‘RPL27A’, ‘RPL28’, ‘RPL3’, ‘RPL31’, ‘RPL32’, ‘RPL34’, ‘RPL37’, ‘RPL37A’, ‘RPL41’, ‘RPL5’, ‘RPL7’, ‘RPL7A’, ‘RPS11’, ‘RPS14’, ‘RPS15A’, ‘RPS16’, ‘RPS19’, ‘RPS2’, ‘RPS20’, ‘RPS23’, ‘RPS24’, ‘RPS27’, ‘RPS3’, ‘RPS3A’, ‘RPS5’, ‘RPS9’, ‘RPSA’, ‘SEC11A’, ‘SLC25A3’, ‘SLC25A5’, ‘SNHG16’, ‘SNHG5’, ‘TOMM20’]. For each gene, we calculated its z score by using the mean and standard deviation from the reference control data. The z score of the target gene program is our measurement, which will be referred to as the score. I can only do a few rounds of experimentation. Your task is predicting and prioritizing genes for the gene panel design in each round to find the most relevant genes (hit genes) to the research problem.

##### Previous Round’s result

This is your 2nd round for predicting new genes. You have 10 rounds in total. So far, the measured scores of tested genes that have not shown effectiveness toward our goal are: translation

DDX5 0.099653 DDX6 -0.439711 … *<* Experiment Results *>*

You have successfully identified 4 hits over all experimental rounds!

Hit genes with positive scores: Empty

Hit genes with negative scores: RIOK2 -1.034148 RPS27A -0.917903 RPS29 -0.165591 RPS28 -0.520588.

In this round, we aim to test 64 genes. Your task is to predict and prioritize genes for the upcoming experiment to find as many hit genes as possible. Before making your predictions, you are encouraged to thoroughly analyze the current data and apply critical thinking.

##### Action Pool

There are several actions you can choose from to learn more about the problem and previous data 1. **Gene Set Enrichment Analysis (GSEA):** Request the GSEA on the control cell. The control cell often bears non-targeting single-guide RNAs (sgRNAs). It serves as a baseline reference in the experiment, providing an unperturbed transcriptomic profile against which the effects of genetic perturbations can be compared.We will return up to the top 10 enriched pathways with p-values smaller than 0.05, based on different libraries.2. **Top Predicted Genes:** Request the top 64 genes most likely to have a high (high predicted absolute value) impact on the target gene program, predicted by an XGBoost model. The model will be trained on previous experimental data, the input of the model is the embedding of one perturbation, and the output is the predicted score of the target gene expression program under the input perturbation.3. **Critical Thinking:** Reflect on the current data and output your analysis, providing insights for further investigation. 5. **Output Prediction:** Indicate that you are ready to output your predictions. You will provide and prioritize genes following a specific format. Call this action when your prediction does not exceed the number of genes you need to predict. If you want to refine your existing predictions, choose action 9.6. **Gene Pathway Enrichment Analysis on Positive Hit Genes:** Request the gene pathway enrichment analysis on hit genes with positive scores. We will return up to the top 10 enriched pathways with p-values smaller than 0.05, based on different libraries. And we will provide the most frequent valid genes shared in these enriched pathways.7. **Gene Pathway Enrichment Analysis on Negative Hit Genes:** Request the gene pathway enrichment analysis on hit genes with negative scores. We will return up to the top 10 enriched pathways with p-values smaller than 0.05, based on different libraries. And we will provide the most frequent valid genes shared in these enriched pathways.9. **Prediction Refinement:** Refine your prediction based on new observations and feedback.10. **Cut-off & Complete:** Finish the current round and output the final prediction. If the length of the current solution exceeds the number of genes you need to predict, the first 64 genes will be selected as the final prediction.11. **Gene Set Enrichment Analysis (GSEA) on Newly Experimented Hit Genes:** Request the gene set enrichment analysis on hit genes that have been experimented on in the previous round. We will return up to the top 3 enriched pathways with p-values smaller than 0.05 for each hit perturbation, based on the same library.Note that you can choose an option more than once, but each time you can only choose one. In total, you can choose options up to 20 times.All valid actions are 1, 3, 5, 7, 11. Provide only your most desired choice in the format *<*STEP*>*Number*<*STEP*>* without or with no more than two sentences of explanation.

##### Execution Log

**PerTurboAgent**: *<*STEP*>*6*<*STEP*>*

**System**: Your choice is 6. And we performed the enrichement analysis based on three libraries. For the first library Reactome 2022, the top enriched pathway is [‘CD28 Dependent PI3K/Akt Signaling R-HSA-389357’, ‘mTORC1-mediated Signaling R-HSA-166208’, ‘Constitutive Signaling By AKT1 E17K In Cancer R-HSA-5674400’, ‘VEGFR2 Mediated Vascular Permeability R-HSA-5218920’, ‘Energy Dependent Regulation Of mTOR By LKB1-AMPK R-HSA-380972’, ‘CD28 Co-Stimulation R-HSA-389356’, ‘HSF1-dependent Transactivation R-HSA-3371571’, ‘Regulation Of TP53 Degradation R-HSA-6804757’, ‘Regulation Of TP53 Expression And Degradation R-HSA-6806003’, ‘MTOR Signaling R-HSA-165159’]. For the second library MSigDB Hallmark 2020, the top enriched pathway is [‘Spermatogenesis’]. For the third library GO Biological Process 2023, the top enriched pathway is [‘Regulation Of Pentose-Phosphate Shunt (GO:0043456)’, ‘Regulation Of Wound Healing, Spreading Of Epidermal Cells (GO:1903689)’, ‘Regulation Of Myeloid Leukocyte Differentiation (GO:0002761)’, ‘Anoikis (GO:0043276)’, ‘Positive Regulation Of Keratinocyte Migration (GO:0051549)’, ‘Positive Regulation Of Transcription Of Nucleolar Large rRNA By RNA Polymerase I (GO:1901838)’, ‘Response To Leucine (GO:0043201)’, ‘Cellular Response To Leucine (GO:0071233)’, ‘TORC1 Signaling (GO:0038202)’, ‘Regulation Of Keratinocyte Migration (GO:0051547)’]. These pathways span various biological processes and maybe linked to different diseases or cellular function. The gene names provided here have been identified as the candidates for most frequently occurring genes across all enriched pathway: [‘MLST8’, ‘RPTOR’, ‘PDPK1’, ‘PRR5’, ‘RICTOR’, ‘MAPKAP1’, ‘RRAGD’, ‘LAMTOR1’, ‘LAMTOR4’, ‘AKT1S1’, ‘LAMTOR3’, ‘LAMTOR2’, ‘LAMTOR5’, ‘TRIB3’, ‘THEM4’, ‘HOXC13’, ‘SLC38A9’, ‘RRAGA’, ‘RHEB’, ‘RRAGC’, ‘RRAGB’, ‘MDM2’, ‘STK11’, ‘UBB’, ‘CDK1’, ‘PIK3R2’, ‘MAP3K8’, ‘PIK3R1’, ‘FYN’, ‘FKBP1A’, ‘RPS6KB1’, ‘EIF4EBP1’, ‘YWHAB’, ‘EIF4G1’, ‘RPS6’, ‘EEF2K’, ‘EIF4E’, ‘EIF4B’, ‘RNASE1’, ‘VAV1’, ‘HSP90AA1’, ‘PKN1’, ‘ARHGEF7’, ‘PKN2’, ‘STRADA’, ‘CAB39L’, ‘PRKAG1’, ‘PRKAA1’, ‘CAB39’, ‘TSC1’, ‘PRKAB1’, ‘PRKAB2’, ‘STRADB’, ‘PPM1A’, ‘DAXX’, ‘CCNG1’, ‘PPP2R5C’, ‘SGK1’, ‘PPP2R1A’, ‘RNF34’, ‘UBA52’, ‘PHF20’, ‘PPP2CA’, ‘CDK2’]

**System**: This is your 2/20 time to take action. Based on the current observations, Provide the number of your choice with format *<*STEP*>*Number*<*/STEP*>* and I will provide the information accordingly.

**PerTurboAgent**: *<*STEP*>*11*<*/STEP*>*

**System**: Your choice is 11. And we performed the enrichement analysis on every new experimented perturbation based on the library Reactome 2022.For the perturbation MTOR, the top enriched pathway is [‘rRNA Processing R-HSA-72312’, ‘rRNA Processing In Nucleus And Cytosol R-HSA-8868773’, ‘Major Pathway Of rRNA Processing In Nucleolus And Cytosol R-HSA-6791226’]”,

**System**: This is your 3/20 time to take action. Based on the current observations, Provide the number of your choice with format *<*STEP*>*Number*<*/STEP*>* and I will provide the information accordingly

**PerTurboAgent**: *<*STEP*>*3*<*/STEP*>*

**System**: Your choice is 3. Show your reflections here.

**PerTurboAgent**: ‘Based on the enrichment analyses, both MTOR and the new hit genes from positive scores are heavily involved in signaling pathways related to mTOR and Akt signaling, which are crucial for translation control and cell growth. The presence of pathways like “rRNA Processing” in the MTOR perturbation reinforces the notion that targeting translation regulation is a viable strategy. The enriched pathways from the previous hit gene (MTOR) suggest the involvement of processes closely associated with translation and ribosome biogenesis, which aligns with the goal of identifying perturbations affecting translation-related gene programs. For the next round of predictions, considering genes associated with the enriched pathways and processes identified could yield valuable insights. Prioritizing genes involved in “mTOR Signaling”, “TORC1 Signaling”, and rRNA processing pathways might enhance the likelihood of identifying additional hit genes.’,

**System**: This is your 4/20 time to take action. Based on the current observations, Provide the number of your choice with format *<*STEP*>*Number*<*/STEP*>* and I will provide the information accordingly

**PerTurboAgent**: *<*STEP*>*1*<*/STEP*>*

**System**: Your choice is 1. And we performed the enrichement analysis based on three libraries. For the first library MSigDB Hallmark 2020, the top enriched pathway is [‘Myc Targets V1’, ‘Allograft Rejection’]. For the second library Reactome 2022, the top enriched pathway is [‘Scavenging By Class A Receptors R-HSA-3000480’, ‘Modulation By Mtb Of Host Immune System R-HSA-9637628’, ‘Uptake And Function Of Diphtheria Toxin R-HSA-5336415’, ‘FLT3 Signaling By CBL Mutants R-HSA-9706377’, ‘Myoclonic Epilepsy Of Lafora R-HSA-3785653’, ‘TFAP2A Acts As A Transcriptional Repressor During Retinoic Acid Induced Cell Differentiation R-HSA-8869496’, ‘Noncanonical Activation Of NOTCH3 R-HSA-9017802’, ‘rRNA Processing R-HSA-72312’, ‘Glycogen Synthesis R-HSA-3322077’, ‘Downregulation Of ERBB4 Signaling R-HSA-1253288’]. For the third library KEGG 2021 Human, the top enriched pathway is [‘Ribosome’, ‘Coronavirus disease’, ‘Type I diabetes mellitus’, ‘Phototransduction’, ‘Systemic lupus erythematosus’, ‘Parkinson disease’, ‘Proteasome’, ‘Oxidative phosphorylation’, ‘Mineral absorption’, ‘Arachidonic acid metabolism’]. These pathways span various biological processes and maybe linked to different diseases or cellular function. The gene names provided here have been identified as the candidates for most frequently occurring genes across all enriched pathway: [‘RPS27A’, ‘UBA52’, ‘UBB’, ‘UBC’, ‘RPL34’, ‘RPL22’, ‘RPLP0’, ‘RPS10’, ‘RPL18’, ‘RPL6’, ‘RPS2’, ‘RPS5’, ‘RPS3’, ‘RPS6’, ‘RPL14’, ‘RPL9’, ‘RPS3A’, ‘RPL39’, ‘RPS19’, ‘RPS9’, ‘PSMD1’, ‘PSMC4’, ‘CYC1’, ‘PSMA1’, ‘HSPD1’, ‘PSMD7’, ‘PSMD3’, ‘NDUFAB1’, ‘PSMD14’, ‘PSMA6’, ‘PSMA2’, ‘PSMA4’, ‘NPM1’, ‘PSMD8’, ‘COX5A’, ‘PSMA7’, ‘PSMB2’, ‘PSMB3’, ‘PSMC6’, ‘RPS14’, ‘RPS8’, ‘RPS29’, ‘RPL26L1’, ‘RPS4X’, ‘RPL21’, ‘RPL37A’, ‘RPL27’, ‘RPS27L’, ‘RPL41’, ‘RPL28’, ‘RPS21’, ‘RPL36’, ‘RPL10A’, ‘RPLP2’, ‘RPL19’, ‘RPL37’, ‘RPS12’, ‘RPL31’, ‘RPL26’, ‘RPL29’, ‘RPL8’, ‘RPS11’, ‘RPL32’, ‘RPS28’]”,

**System**: This is your 5/20 time to take action. Based on the current observations, Provide the number of your choice with format *<*STEP*>*Number*<*/STEP*>* and I will provide the information accordingly

**PerTurboAgent**: *<*STEP*>*5*<*/STEP*>*

**System**: Your choice is 5. You have to predict 64 genes. Use HGNC gene naming convention. DO NOT PREDICT GENES THAT HAVE ALREADY BEEN TESTED OR PREDICTED.Respond in this format exactly:1. Reflection: Thoughts on previous results and next steps. 2. Research Plan: The full high level research plan, with current status and reasoning behind each proposed approach. It should be at most 5 sentences.3. Solution: A list of predicted genes: [*<*Gene name 1*>, <*Gene name 2*>*, …]

**PerTurboAgent**: 1. Reflection: The enrichment analyses have highlighted critical pathways involving mTOR signaling, translation-related processes, and ribosomal biogenesis. The convergence of these pathways suggests that targeting genes involved in these processes could lead to successful identification of hit genes affecting translation-related gene programs. Furthermore, the gene enrichment from the control perturbations suggests potential novel markers and pathways unexplored in previous rounds.2. Research Plan: We aim to prioritize genes related to the mTOR, ribosomal, and Akt pathways based on previous enrichment analyses and pathways linked to translation control. The current status reveals one hit gene linked to the mTOR pathway and suggests focusing on genes associated with translation and ribosome-related pathways. We will leverage insights from known interactions within these pathways and exclude previously tested genes to maximize candidate diversity. Our objective is to increase the identification of impactful genes across successive rounds, refining gene selection iteratively.3. Solution: [‘MLST8’, ‘RPTOR’, ‘PDPK1’, ‘PRR5’, ‘RICTOR’, ‘MAPKAP1’, ‘RRAGD’, ‘LAMTOR1’, ‘LAMTOR4’, ‘AKT1S1’, ‘LAMTOR3’, ‘LAMTOR2’, ‘LAMTOR5’, ‘TRIB3’, ‘THEM4’, ‘HOXC13’, ‘SLC38A9’, ‘RRAGA’, ‘RHEB’, ‘RRAGC’, ‘RRAGB’, ‘MDM2’, ‘STK11’, ‘CDK1’, ‘PIK3R2’, ‘MAP3K8’, ‘PIK3R1’, ‘FYN’, ‘FKBP1A’, ‘RPS6KB1’, ‘EIF4EBP1’, ‘YWHAB’, ‘EIF4G1’, ‘EEF2K’, ‘EIF4E’, ‘EIF4B’, ‘RNASE1’, ‘VAV1’, ‘HSP90AA1’, ‘PKN1’, ‘ARHGEF7’, ‘PKN2’, ‘STRADA’, ‘CAB39L’, ‘PRKAG1’, ‘PRKAA1’, ‘CAB39’, ‘TSC1’, ‘PRKAB1’, ‘PRKAB2’, ‘STRADB’, ‘PPM1A’, ‘DAXX’, ‘CCNG1’, ‘PPP2R5C’, ‘SGK1’, ‘PPP2R1A’, ‘RNF34’, ‘PHF20’, ‘PPP2CA’, ‘RPS27A’, ‘UBC’, ‘UBA52’, ‘RPL34’]

**System**: Now, there are 64 valid genes: [‘MLST8’, ‘RPTOR’, ‘PDPK1’, ‘PRR5’, ‘RICTOR’, ‘MAPKAP1’, ‘RRAGD’, ‘LAMTOR1’, ‘LAMTOR4’, ‘AKT1S1’, ‘LAMTOR3’, ‘LAMTOR2’, ‘LAMTOR5’, ‘TRIB3’, ‘THEM4’, ‘HOXC13’, ‘SLC38A9’, ‘RRAGA’, ‘RHEB’, ‘RRAGC’, ‘RRAGB’, ‘MDM2’, ‘STK11’, ‘CDK1’, ‘PIK3R2’, ‘MAP3K8’, ‘PIK3R1’, ‘FYN’, ‘FKBP1A’, ‘RPS6KB1’, ‘EIF4EBP1’, ‘YWHAB’, ‘EIF4G1’, ‘EEF2K’, ‘EIF4E’, ‘EIF4B’, ‘RNASE1’, ‘VAV1’, ‘HSP90AA1’, ‘PKN1’, ‘ARHGEF7’, ‘PKN2’, ‘STRADA’, ‘CAB39L’, ‘PRKAG1’, ‘PRKAA1’, ‘CAB39’, ‘TSC1’, ‘PRKAB1’, ‘PRKAB2’, ‘STRADB’, ‘PPM1A’, ‘DAXX’, ‘CCNG1’, ‘PPP2R5C’, ‘SGK1’, ‘PPP2R1A’, ‘RNF34’, ‘PHF20’, ‘PPP2CA’, ‘RPS27A’, ‘UBC’, ‘UBA52’, ‘RPL34’]. There are 0 invalid or duplicated genes: []

**System**: This is your 6/20 time to take action. Based on the current observations, Provide the number of your choice with format *<*STEP*>*Number*<*/STEP*>* and I will provide the information accordingly.

**PerTurboAgent**: *<*STEP*>*3*<*/STEP*>*

**System**: Your choice is 3. Show your reflections here.

**PerTurboAgent**: The selected gene list effectively targets key components of the mTOR and ribosomal signaling pathways, which are central to the regulation of translation processes. Many of these genes, such as RPTOR, MLST8, and AKT1S1, are directly involved in the mTORC1 complex, a critical regulator of protein synthesis and cell growth. The inclusion of genes associated with ribosome function and biogenesis, such as RPL34 and RPS27A, complements the research objective of affecting translation-related gene programs. This gene panel, thus, integrates insights from previous enrichment analyses, focusing on genes with potential high impact on the translation process, while avoiding prior tested candidates to diversify experimental outcomes. The current predictions maintain a balance between pathway-centric exploration and novel candidate testing.

**System**: This is your 7/20 time to take action. Based on the current observations, Provide the number of your choice with format *<*STEP*>*Number*<*/STEP*>* and I will provide the information accordingly.

**PerTurboAgent**: *<*STEP*>*6*<*/STEP*>*

**System**: Your choice is 6. And we performed the enrichement analysis based on three libraries. For the first library Reactome 2022, the top enriched pathway is [‘CD28 Dependent PI3K/Akt Signaling R-HSA-389357’, ‘mTORC1-mediated Signaling R-HSA-166208’, ‘Constitutive Signaling By AKT1 E17K In Cancer R-HSA-5674400’, ‘VEGFR2 Mediated Vascular Permeability R-HSA-5218920’, ‘Energy Dependent Regulation Of mTOR By LKB1-AMPK R-HSA-380972’, ‘CD28 Co-Stimulation R-HSA-389356’, ‘HSF1-dependent Transactivation R-HSA-3371571’, ‘Regulation Of TP53 Degradation R-HSA-6804757’, ‘Regulation Of TP53 Expression And Degradation R-HSA-6806003’, ‘MTOR Signaling R-HSA-165159’]. For the second library KEGG 2021 Human, the top enriched pathway is [‘Type II diabetes mellitus’, ‘Acute myeloid leukemia’, ‘Adipocytokine signaling pathway’, ‘Central carbon metabolism in cancer’, ‘Glioma’, ‘Pancreatic cancer’, ‘ErbB signaling pathway’, ‘Colorectal cancer’, ‘PD-L1 expression and PD-1 checkpoint pathway in cancer’, ‘Prostate cancer’]. For the third library GO Biological Process 2023, the top enriched pathway is [‘Regulation Of Pentose-Phosphate Shunt (GO:0043456)’, ‘Regulation Of Wound Healing, Spreading Of Epidermal Cells (GO:1903689)’, ‘Regulation Of Myeloid Leukocyte Differentiation (GO:0002761)’, ‘Anoikis (GO:0043276)’, ‘Positive Regulation Of Keratinocyte Migration (GO:0051549)’, ‘Positive Regulation Of Transcription Of Nucleolar Large rRNA By RNA Polymerase I (GO:1901838)’, ‘Response To Leucine (GO:0043201)’, ‘Cellular Response To Leucine (GO:0071233)’, ‘TORC1 Signaling (GO:0038202)’, ‘Regulation Of Keratinocyte Migration (GO:0051547)’]. These pathways span various biological processes and maybe linked to different diseases or cellular function. The gene names provided here have been identified as the candidates for most frequently occurring genes across all enriched pathway: [‘PIK3CD’, ‘MAPK1’, ‘PIK3CB’, ‘MAPK3’, ‘NRAS’, ‘RPS6KB2’, ‘CHUK’, ‘BAD’, ‘GRB2’, ‘IKBKB’, ‘IKBKG’, ‘PTEN’, ‘MAPK9’, ‘MAPK8’, ‘RELA’, ‘SOS1’, ‘NFKB1’, ‘GSK3B’, ‘CASP9’, ‘STAT3’, ‘CDKN1B’, ‘UBB’, ‘CAMK2G’, ‘CCNA2’, ‘CDK2’, ‘TCF7’, ‘TCF7L2’, ‘TCF7L1’, ‘LEF1’, ‘NFKBIA’, ‘DDB2’, ‘E2F3’, ‘POLK’, ‘BAK1’, ‘PLCG1’, ‘RB1’, ‘E2F2’, ‘GADD45A’, ‘JUN’, ‘RPS6’, ‘CREB1’, ‘FOXO1’, ‘CALM1’, ‘JUP’, ‘CDC42’, ‘HSP90AB1’, ‘PPP2R1B’, ‘PPP2CB’, ‘MDM4’, ‘USP2’, ‘RFFL’, ‘USP7’, ‘HK1’, ‘SOCS3’, ‘HK2’, ‘PKM’, ‘STAT5B’, ‘STAT5A’, ‘NFKBIE’, ‘NFKBIB’, ‘SLC2A1’, ‘PTPN11’, ‘PRKCQ’, ‘HIF1A’]

**System**: This is your 8/20 time to take action. Based on the current observations, Provide the number of your choice with format *<*STEP*>*Number*<*/STEP*>* and I will provide the information accordingly.

**PerTurboAgent**: *<*STEP*>*3*<*/STEP*>*

**System**: Your choice is 3. Show your reflections here.

**PerTurboAgent**: The pathway enrichment analysis indicates a strong association between the identified genes and pathways linked to mTOR signaling, AKT signaling, translation regulation, and cancer-related processes like cell survival and proliferation. The presence of pathways such as “mTORC1-mediated Signaling” and “TORC1 Signaling” reiterates their importance in regulating protein synthesis and cellular growth, aligning well with the goal of impacting translation-related gene programs. Additionally, the connection to diseases such as diabetes and cancers further highlights these pathwaysśystemic importance. This reinforces prioritizing genes interacting within these pathways for subsequent rounds, as their perturbation could unveil significant impacts on the translation program, providing deeper insights into their biological roles.

**System**: This is your 9/20 time to take action. Based on the current observations, Provide the number of your choice with format *<*STEP*>*Number*<*/STEP*>* and I will provide the information accordingly.

PerTurboAgent: *<*STEP*>*9*<*/STEP*>*

**System**: Your choice is 9. Your current prediction has 64 genes. Think carefully, critique the current plan and propose the genes you want to remove and add. All other genes will be kept. Use HGNC gene naming convention. DO NOT PREDICT GENES THAT HAVE ALREADY BEEN TESTED.Please do not make changes if there is no need to make a change. Respond in this format exactly:1. Critique: include all relevant details of the critique.2. Updated_Solution_Removal: Give a list of genes you want to remove from the current prediction separated by commas in this format: [*<*Gene name 1*>, <*Gene name 2*>*, …]3. Updated_Solution_Addition: Give a list of genes you want to add to the current prediction separated by commas in this format: [*<*Gene name 1*>, <*Gene name 2*>*, …]

**PerTurboAgent**: 1. Critique: The current prediction effectively captures key genes linked to translation-related pathways, but can further be refined by leveraging the latest enrichment analysis. This analysis highlighted additional genes involved in pathways relevant to translation and cell growth, not previously considered. Incorporating genes such as PIK3CD and MAPK1, which are integral to the enriched signaling pathways, may provide more comprehensive insights. Therefore, it’s beneficial to replace genes with less direct involvement in these core pathways to achieve an optimized gene set for testing.2. Updated_Solution_Removal: [‘HOXC13’, ‘RNASE1’, ‘CAB39L’, ‘CAB39’]3. Updated_Solution_Addition: [‘PIK3CD’, ‘MAPK1’, ‘GRB2’, ‘PTEN’]

**System**: Now, there are 64 valid genes: [‘MLST8’, ‘RPTOR’, ‘PDPK1’, ‘PRR5’, ‘RICTOR’, ‘MAPKAP1’, ‘RRAGD’, ‘LAMTOR1’, ‘LAMTOR4’, ‘AKT1S1’, ‘LAMTOR3’, ‘LAMTOR2’, ‘LAMTOR5’, ‘TRIB3’, ‘THEM4’, ‘SLC38A9’, ‘RRAGA’, ‘RHEB’, ‘RRAGC’, ‘RRAGB’, ‘MDM2’, ‘STK11’, ‘CDK1’, ‘PIK3R2’, ‘MAP3K8’, ‘PIK3R1’, ‘FYN’, ‘FKBP1A’, ‘RPS6KB1’, ‘EIF4EBP1’, ‘YWHAB’, ‘EIF4G1’, ‘EEF2K’, ‘EIF4E’, ‘EIF4B’, ‘VAV1’, ‘HSP90AA1’, ‘PKN1’, ‘ARHGEF7’, ‘PKN2’, ‘STRADA’, ‘PRKAG1’, ‘PRKAA1’, ‘TSC1’, ‘PRKAB1’, ‘PRKAB2’, ‘STRADB’, ‘PPM1A’, ‘DAXX’, ‘CCNG1’, ‘PPP2R5C’, ‘SGK1’, ‘PPP2R1A’, ‘RNF34’, ‘PHF20’, ‘PPP2CA’, ‘RPS27A’, ‘UBC’, ‘UBA52’, ‘RPL34’, ‘PIK3CD’, ‘MAPK1’, ‘GRB2’, ‘PTEN’]. There are 0 invalid or duplicated genes: [].

## References

Josh Achiam, Steven Adler, Sandhini Agarwal, Lama Ahmad, Ilge Akkaya, Florencia Leoni Aleman, Diogo Almeida, Janko Altenschmidt, Sam Altman, Shyamal Anadkat, et al. Gpt-4 technical report. arXiv preprint 2303.08774, 2023.

Britt Adamson, Thomas M Norman, Marco Jost, Min Y Cho, James K Nuñez, Yuwen Chen, Jacqueline E Villalta, Luke A Gilbert, Max A Horlbeck, Marco Y Hein, et al. A multiplexed single-cell crispr screening platform enables systematic dissection of the unfolded protein response. Cell, 167(7):1867–1882, 2016.

Jean-Philippe Brunet, Pablo Tamayo, Todd R Golub, and Jill P Mesirov. Metagenes and molecular pattern discovery using matrix factorization. Proceedings of the national academy of sciences, 101(12):4164–4169, 2004.

Charlotte Bunne, Stefan G Stark, Gabriele Gut, Jacobo Sarabia Del Castillo, Mitch Levesque, Kjong-Van Lehmann, Lucas Pelkmans, Andreas Krause, and Gunnar Rätsch. Learning single-cell perturbation responses using neural optimal transport. Nature methods, 20(11):1759–1768, 2023.

Yiqun Chen and James Zou. Genept: a simple but effective foundation model for genes and cells built from chatgpt. bioRxiv, pp. 2023–10, 2024.

Haotian Cui, Chloe Wang, Hassaan Maan, Kuan Pang, Fengning Luo, Nan Duan, and Bo Wang. scgpt: toward building a foundation model for single-cell multi-omics using generative ai. Nature Methods, pp. 1–11, 2024.

Atray Dixit, Oren Parnas, Biyu Li, Jenny Chen, Charles P Fulco, Livnat Jerby-Arnon, Nemanja D Marjanovic, Danielle Dionne, Tyler Burks, Raktima Raychowdhury, et al. Perturb-seq: dissecting molecular circuits with scalable single-cell rna profiling of pooled genetic screens. cell, 167(7):1853–1866, 2016.

Abhimanyu Dubey, Abhinav Jauhri, Abhinav Pandey, Abhishek Kadian, Ahmad Al-Dahle, Aiesha Letman, Akhil Mathur, Alan Schelten, Amy Yang, Angela Fan, et al. The llama 3 herd of models. arXiv preprint 2407.21783, 2024.

Shanghua Gao, Ada Fang, Yepeng Huang, Valentina Giunchiglia, Ayush Noori, Jonathan Richard Schwarz, Yasha Ektefaie, Jovana Kondic, and Marinka Zitnik. Empowering biomedical discovery with ai agents. Cell, 187(22):6125–6151, 2024.

Kathryn Geiger-Schuller, Basak Eraslan, Olena Kuksenko, Kushal K Dey, Karthik A Jagadeesh, Pratiksha I Thakore, Ozge Karayel, Andrea R Yung, Anugraha Rajagopalan, Ana M Meireles, et al. Systematically characterizing the roles of e3-ligase family members in inflammatory responses with massively parallel perturb-seq. bioRxiv, pp. 2023–01, 2023.

Madalina Giurgiu, Julian Reinhard, Barbara Brauner, Irmtraud Dunger-Kaltenbach, Gisela Fobo, Goar Frishman, Corinna Montrone, and Andreas Ruepp. Corum: the comprehensive resource of mammalian protein complexes—2019. Nucleic acids research, 47(D1):D559–D563, 2019.

Stewart Greenhill, Santu Rana, Sunil Gupta, Pratibha Vellanki, and Svetha Venkatesh. Bayesian optimization for adaptive experimental design: A review. IEEE access, 8:13937–13948, 2020.

Minsheng Hao, Jing Gong, Xin Zeng, Chiming Liu, Yucheng Guo, Xingyi Cheng, Taifeng Wang, Jianzhu Ma, Xuegong Zhang, and Le Song. Large-scale foundation model on single-cell transcriptomics. Nature Methods, pp. 1–11, 2024.

Kaixuan Huang, Yuanhao Qu, Henry Cousins, William A Johnson, Di Yin, Mihir Shah, Denny Zhou, Russ Altman, Mengdi Wang, and Le Cong. Crispr-gpt: An llm agent for automated design of gene-editing experiments. arXiv preprint 2404.18021, 2024a.

Kexin Huang, Romain Lopez, Jan-Christian Hütter, Takamasa Kudo, Antonio Rios, and Aviv Regev. Sequential optimal experimental design of perturbation screens guided by multi-modal priors. In International Conference on Research in Computational Molecular Biology, pp. 17–37. Springer, 2024b.

Guolin Ke, Qi Meng, Thomas Finley, Taifeng Wang, Wei Chen, Weidong Ma, Qiwei Ye, and Tie-Yan Liu. Lightgbm: A highly efficient gradient boosting decision tree. Advances in neural information processing systems, 30, 2017.

Alexey Kurakin, Ian J Goodfellow, and Samy Bengio. Adversarial examples in the physical world. In Artificial intelligence safety and security, pp. 99–112. Chapman and Hall/CRC, 2018.

Woosuk Kwon, Zhuohan Li, Siyuan Zhuang, Ying Sheng, Lianmin Zheng, Cody Hao Yu, Joseph E. Gonzalez, Hao Zhang, and Ion Stoica. Efficient memory management for large language model serving with pagedattention. In Proceedings of the ACM SIGOPS 29th Symposium on Operating Systems Principles, 2023.

Alexander Lachmann, Zhuorui Xie, and Avi Ma’ayan. blitzgsea: efficient computation of gene set enrichment analysis through gamma distribution approximation. Bioinformatics, 38(8):2356– 2357, 2022.

Yongju Lee, Dyke Ferber, Jennifer E Rood, Aviv Regev, and Jakob Nikolas Kather. How ai agents will change cancer research and oncology. Nature Cancer, pp. 1–3, 2024.

Mohammad Lotfollahi, F Alexander Wolf, and Fabian J Theis. scgen predicts single-cell perturbation responses. Nature methods, 16(8):715–721, 2019.

Mohammad Lotfollahi, Anna Klimovskaia Susmelj, Carlo De Donno, Leon Hetzel, Yuge Ji, Ignacio L Ibarra, Sanjay R Srivatsan, Mohsen Naghipourfar, Riza M Daza, Beth Martin, et al. Predicting cellular responses to complex perturbations in high-throughput screens. Molecular systems biology, 19(6):e11517, 2023.

Chris Lu, Cong Lu, Robert Tjarko Lange, Jakob Foerster, Jeff Clune, and David Ha. The ai scientist: Towards fully automated open-ended scientific discovery. arXiv preprint 2408.06292, 2024.

Pan Lu, Swaroop Mishra, Tony Xia, Liang Qiu, Kai-Wei Chang, Song-Chun Zhu, Oyvind Tafjord, Peter Clark, and Ashwin Kalyan. Learn to explain: Multimodal reasoning via thought chains for science question answering. In The 36th Conference on Neural Information Processing Systems (NeurIPS), 2022.

Laura Luebbert and Lior Pachter. Efficient querying of genomic reference databases with gget. Bioinformatics, 39(1):btac836, 2023.

Clare Lyle, Arash Mehrjou, Pascal Notin, Andrew Jesson, Stefan Bauer, Yarin Gal, and Patrick Schwab. Discobax discovery of optimal intervention sets in genomic experiment design. In International Conference on Machine Learning, pp. 23170–23189. PMLR, 2023.

Arash Mehrjou, Ashkan Soleymani, Andrew Jesson, Pascal Notin, Yarin Gal, Stefan Bauer, and Patrick Schwab. Genedisco: A benchmark for experimental design in drug discovery. arXiv preprint 2110.11875, 2021.

Djordje Miladinovic, Tobias Höppe, Mathieu Chevalley, Andreas Georgiou, Lachlan Stuart, Arash Mehrjou, Marcus Bantscheff, Bernhard Schölkopf, and Patrick Schwab. In-silico biological discovery with large perturbation models. arXiv preprint 2503.23535, 2025.

Ajay Nadig, Joseph M Replogle, Angela N Pogson, Steven A McCarroll, Jonathan S Weissman, Elise B Robinson, and Luke J O’Connor. Transcriptome-wide characterization of genetic perturbations. bioRxiv, 2024.

Thomas M Norman, Max A Horlbeck, Joseph M Replogle, Alex Y Ge, Albert Xu, Marco Jost, Luke A Gilbert, and Jonathan S Weissman. Exploring genetic interaction manifolds constructed from rich single-cell phenotypes. Science, 365(6455):786–793, 2019.

Long Ouyang, Jeffrey Wu, Xu Jiang, Diogo Almeida, Carroll Wainwright, Pamela Mishkin, Chong Zhang, Sandhini Agarwal, Katarina Slama, Alex Ray, et al. Training language models to follow instructions with human feedback. Advances in neural information processing systems, 35:27730–27744, 2022.

Fabian Pedregosa, Gaël Varoquaux, Alexandre Gramfort, Vincent Michel, Bertrand Thirion, Olivier Grisel, Mathieu Blondel, Peter Prettenhofer, Ron Weiss, Vincent Dubourg, et al. Scikit-learn: Machine learning in python. the Journal of machine Learning research, 12:2825–2830, 2011.

Rafael Rafailov, Archit Sharma, Eric Mitchell, Christopher D Manning, Stefano Ermon, and Chelsea Finn. Direct preference optimization: Your language model is secretly a reward model. Advances in Neural Information Processing Systems, 36:53728–53741, 2023.

Joseph M Replogle, Reuben A Saunders, Angela N Pogson, Jeffrey A Hussmann, Alexander Lenail, Alina Guna, Lauren Mascibroda, Eric J Wagner, Karen Adelman, Gila Lithwick-Yanai, et al. Mapping information-rich genotype-phenotype landscapes with genome-scale perturb-seq. Cell, 185(14):2559–2575, 2022.

Jennifer E Rood, Anna Hupalowska, and Aviv Regev. Toward a foundation model of causal cell and tissue biology with a perturbation cell and tissue atlas. Cell, 187(17):4520–4545, 2024.

Yusuf Roohani, Kexin Huang, and Jure Leskovec. Predicting transcriptional outcomes of novel multigene perturbations with gears. Nature Biotechnology, 42(6):927–935, 2024a.

Yusuf Roohani, Andrew Lee, Qian Huang, Jian Vora, Zachary Steinhart, Kexin Huang, Alexander Marson, Percy Liang, and Jure Leskovec. Biodiscoveryagent: An ai agent for designing genetic perturbation experiments. arXiv preprint 2405.17631, 2024b.

Ozan Sener and Silvio Savarese. Active learning for convolutional neural networks: A core-set approach. arXiv preprint 1708.00489, 2017.

Kyle Swanson, Wesley Wu, Nash L Bulaong, John E Pak, and James Zou. The virtual lab: Ai agents design new sars-cov-2 nanobodies with experimental validation. bioRxiv, pp. 2024–11, 2024.

Damian Szklarczyk, Annika L Gable, Katerina C Nastou, David Lyon, Rebecca Kirsch, Sampo Pyysalo, Nadezhda T Doncheva, Marc Legeay, Tao Fang, Peer Bork, et al. The string database in 2021: customizable protein–protein networks, and functional characterization of user-uploaded gene/measurement sets. Nucleic acids research, 49(D1):D605–D612, 2021.

Kevin Xia, Kai-Zhan Lee, Yoshua Bengio, and Elias Bareinboim. The causal-neural connection: Expressiveness, learnability, and inference. Advances in Neural Information Processing Systems, 34:10823–10836, 2021.

An Yang, Baosong Yang, Beichen Zhang, Binyuan Hui, Bo Zheng, Bowen Yu, Chengyuan Li, Dayiheng Liu, Fei Huang, Haoran Wei, et al. Qwen2. 5 technical report. arXiv preprint 2412.15115, 2024.

Douglas Yao, Loic Binan, Jon Bezney, Brooke Simonton, Jahanara Freedman, Chris J Frangieh, Kushal Dey, Kathryn Geiger-Schuller, Basak Eraslan, Alexander Gusev, et al. Scalable genetic screening for regulatory circuits using compressed perturb-seq. Nature biotechnology, 42(8):1282–1295, 2024.

Ka Yee Yeung and Walter L. Ruzzo. Principal component analysis for clustering gene expression data. Bioinformatics, 17(9):763–774, 2001.

Xiao Yu, Baolin Peng, Vineeth Vajipey, Hao Cheng, Michel Galley, Jianfeng Gao, and Zhou Yu. Exact: Teaching ai agents to explore with reflective-mcts and exploratory learning. arXiv preprint 2410.02052, 2024.

